# Inflammatory state of lymphatic vessels and miRNA profiles associated with relapse in ovarian cancer patients

**DOI:** 10.1101/2020.02.24.962480

**Authors:** Sarah C Johnson, Sanjukta Chakraborty, Anastasios Drosou, Paula Cunnea, Dimitrios Tzovaras, Katherine Nixon, David C Zawieja, Mariappan Muthuchamy, Christina Fotopoulou, James E Moore

## Abstract

Lymphogenic spread is associated with poor prognosis in epithelial ovarian cancer (EOC), yet little is known regarding roles of non-peri-tumoural lymphatic vessels (LVs) outside the tumour microenvironment that may impact relapse. The aim of this feasibility study was to assess whether inflammatory status of the LVs and/or changes in the miRNA profile of the LVs have potential prognostic and predictive value for overall outcome and risk of relapse. Samples of macroscopically normal human lymph LVs (n=10) were isolated from the external iliac vessels draining the pelvic region of patients undergoing debulking surgery. This was followed by quantification of the inflammatory state (low, medium and high) and presence of cancer-infiltration of each LV using immunohistochemistry. LV miRNA expression profiling was also performed, and analysed in the context of high versus low inflammation, and cancer-infiltrated versus non-cancer-infiltrated. Results were correlated with clinical outcome data including relapse with an average follow-up time of 13.3 months. The presence of a high degree of inflammation correlated significantly with patient relapse (p=0.033). Cancer-infiltrated LVs showed a moderate but non-significant association with relapse (p=0.07). Differential miRNA profiles were identified in cancer-infiltrated LVs and those with high versus low inflammation. In particular, several members of the let-7 family were consistently down-regulated in highly inflamed LVs (>1.8-fold, p<0.05) compared to the less inflamed ones. Down-regulation of the let-7 family appears to be associated with inflammation, but whether inflammation contributes to or is an effect of cancer-infiltration requires further investigation.

## Introduction

Epithelial Ovarian cancer (EOC) accounts for over 4000 deaths in the UK every year [1]. Lymph node (LN) metastases are very common in EOC-patients and associated with a more dismal overall prognosis [2-3]. Total macroscopic tumour clearance with removal of bulky LN is one of the most important prognostic factors for survival post-surgery in EOC [3-4]. There is no therapeutic value in the systematic removal of clinically normal appearing LN in the advanced disease setting, as shown in prospective randomised trials [5]. The involvement and changes in the connecting lymphatic vessels (LVs) and how they are related to the future risk of metastasis and relapse have never been investigated, to our knowledge.

LVs are specialised in the uptake and transport of macromolecules from tissue fluid, consequently permitting lymphogenic spread of cancer cells. Within the tumour microenvironment, intra-tumoural LVs are present, however functional tests in mice have indicated that intra-tumoural lymphatics are non-functional [6,7]. The collapse of intra-tumoural LVs is due to some combination of the high interstitial pressure observed in multiple types of tumours, a lack of lymphatic valves, or induced mechanical pressure of growing tumour cells [6]. However, in both mice and humans, functional peri-tumoural lymphatics remain [6]. LVs can facilitate metastases both passively and through active changes in lymphatic endothelial cell (LEC) expression that aid cancer-cell infiltration, allowing subsequent drainage to LNs [8]. Tumour cells can also arrest within the draining LVs and form in-transit metastases, and in some cases escape the LVs, contributing to loco-regional metastasis, potentially aided by altered LV barrier integrity [8].

Lymphatic endothelial cells (LECs) that line the LVs can also modulate immune cell function, in both anti-tumour immune response and cancer-induced immunosuppression [9]. For example, LEC expression of Protein-Death-Ligand 1 (PD-L1), the ligand for T cell inhibitory receptor, PD-1, can contribute to peripheral tolerance in LNs and tumours [10]. Tumour associated LECs can scavenge and cross-present tumour-antigen to cognate CD8^+^ T cells, displaying the ability to endogenise allergen ovalbumin, then present the exogenous antigen on Major Histocompatibility Complex (MHC) class I molecules, and contribute to inducing tolerance of tumour-specific CD8^+^ T cells [11]. LECs of the LN can acquire peptide MHCII-from dendritic cells and induce tolerance of CD4^+^ T cells via subsequent presentation, in addition to up-regulating MHCII in response to viral-induced inflammation or Interferon-gamma (IFN-γ) [12]. LVs are also sensitive to inflammation-induced signalling molecules which can promote lymphangiogenesis, and induce further secretion of inflammatory molecules that may drain to the LNs [13]. Chronic inflammation in collecting LVs induce changes including LV dilation and inhibition of contractile ability but the effects on metastasis are unknown [14].

MicroRNAs provide vital post-transcriptional regulation of gene expression, a function that is often disrupted by cancer [15]. The ability of a single miRNA to inhibit multiple target genes has led to several studies of altered tumour miRNA expression in ovarian cancer and response to chemotherapy. This includes comparison of miRNA expression in different tumour subsets, in healthy versus diseased ovaries, in patient versus healthy serum, benign versus malignant tumours, expression associated with reduced patient survival, and expression associated with drug-resistance [16]. For example, a study using both EOC cell lines and EOC tumour tissue showed highly elevated miR-221, decreased miR-21 expression, and down-regulation of several members of the let-7 family [17].

Expression of miRNAs in LVs and LECs is altered during inflammation and by nearby tumours. In inflamed LECs, upregulated miR-1236 reduced expression of vascular-endothelial growth factor-3 receptor, inhibiting lymphangiogenesis [18]. A separate study showed that inflamed dermal LECs increased miR-155 expression, a miRNA with known involvement in PD-L1 expression which may contribute to peripheral tolerance changes [19]. Furthermore, a panel of differentially expressed miRNAs was identified in inflamed rat LECs and human LEC culture, indicating effects on inflammation, angiogenesis, endothelial mesenchymal transition, cell proliferation and senescence pathways [13]. Tumour expression profiling has identified changes in lymphangiogenesis-inducing miRNAs, such as miR-126 in oral cancer and miR-128 in non-small cell lung cancer, that could influence metastasis [20,21]. A study using human LEC co-culture models of gastric cancer, representing tumoural or draining LVs, identified changes in differential miRNA expression associated with lymphangiogenesis and with positive lymphatic metastasis [22]. Further miRNA expression studies of draining LVs are lacking but many gene dysregulations were identified in LECs isolated from afferent LVs that drained a metastatic gastric tumour in rats, including up-regulation of chemokine CXCL1 [23].

The aim of this feasibility study was to provide insight into, and assess the prognostic value of the changes in the inflammatory state and miRNA expression of the transporting LVs in EOC. It is expected that these changes will be indicative of the immune response in the LVs and the downstream LN, and therefore could be predictive of patient prognosis.

## Methods

### LV sample collection

LVs were collected from patients undergoing debulking surgery for advanced stage (FIGO IIC, III or IV) epithelial ovarian cancer at Hammersmith Hospital, Imperial College Healthcare NHS Trust, London, United Kingdom (Tables 1 and S1). Informed consent was provided from each patient. Ethics committee approval was obtained from Hammersmith and Queen Charlotte’s and Chelsea Research Ethics Committee (REC reference: 05/Q0406/178), and tissue samples were provided by the Imperial College Healthcare NHS Trust Tissue Bank (ICHTB). The following LV processing steps are summarised in S1 Fig.

**Table 1.**
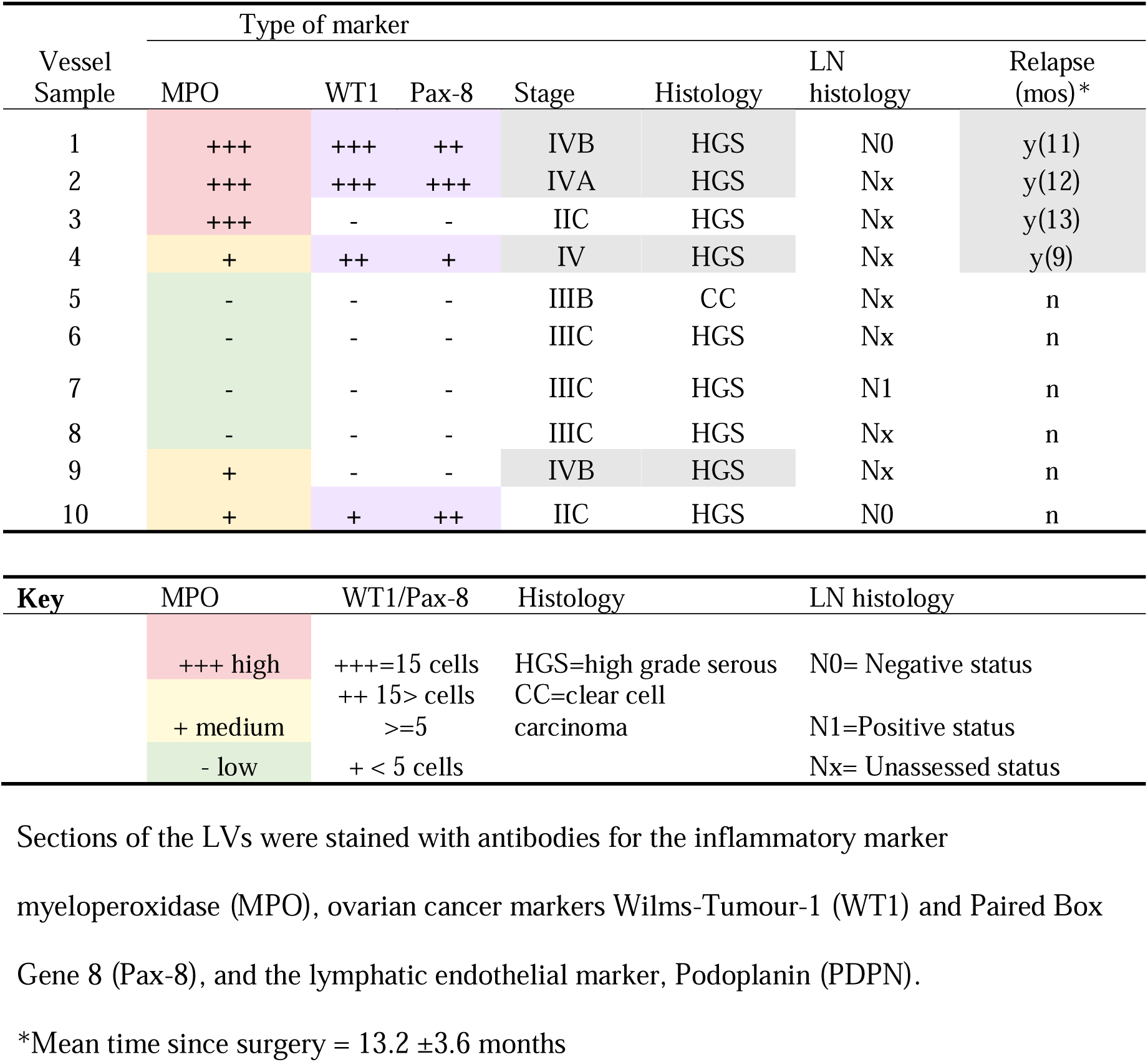
Summary of the LV microscopic state and clinical data.

Macroscopically normal appearing LVs along the external iliac vessels with surrounding fibro-fatty tissue were removed with the nearby LN as part of the debulking procedure. Vessels were ligated with VIcyl03 or PDS-4-0, avoiding bipolar coagulation and other thermic energy to limit vessel damage. All patients had no macroscopic residual disease after surgery.

### Vessel isolation and storage

Tissue specimens were transported directly from surgical theatre to the laboratory in ice-cold sterile Phosphate Buffered Saline (VWR, UK). LVs were identified and isolated along with a small amount of surrounding tissue under a dissection microscope within a sterile laminar flow cabinet (Fig 1 A-C). Each vessel was divided into two sections. One half of each vessel was further cleaned of surrounding fatty tissue, placed in RNA stabilising solution (RNAlater, ThermoFisher, UK), frozen on dry ice and stored at -20°C. The other half was fixed in 4% paraformaldehyde (VWR, UK) for 45 minutes, followed by submersion in 15% sucrose for 3 hours, then 30% sucrose for 6-12 hours. This was followed by embedding in OCT (Scigen, UK) using dry ice and isopropyl alcohol (ThermoFisher, UK) and stored at - 80°C.

**Fig 1.**
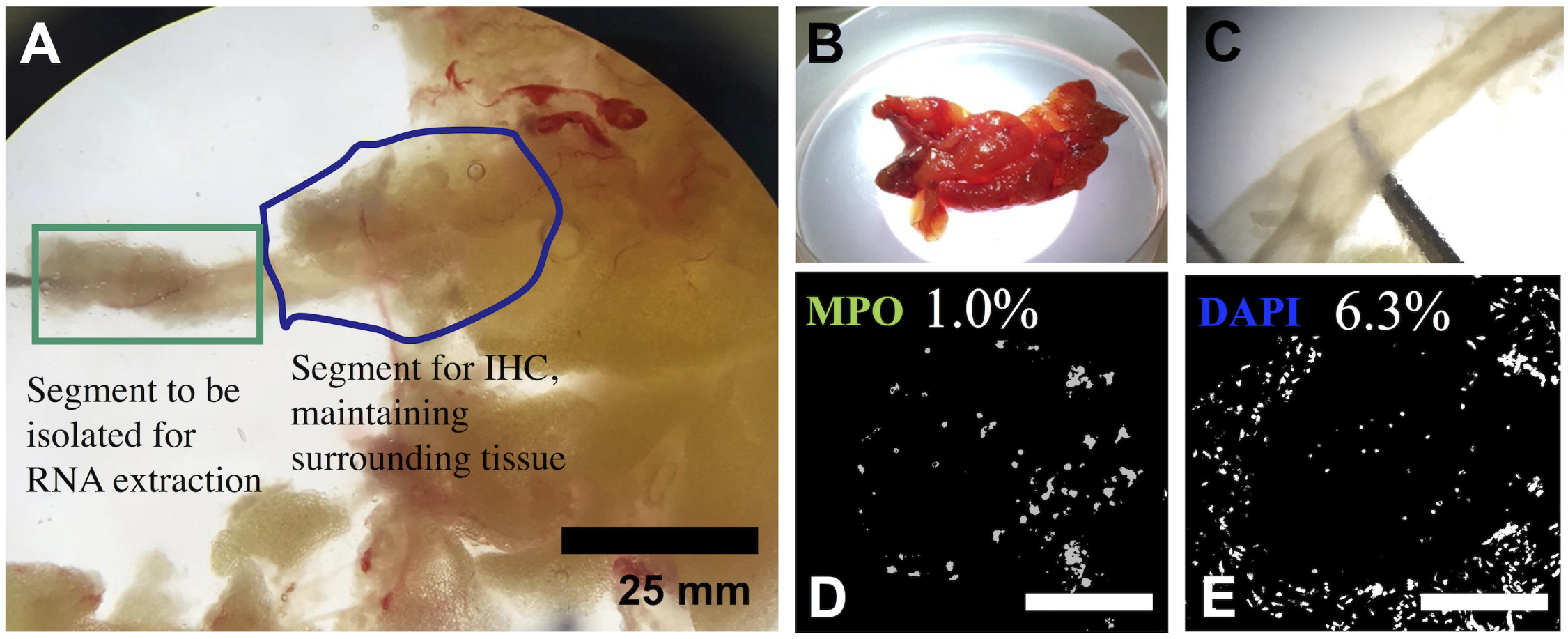
Lymphatic vessel isolation and classification. A. LV isolation technique. B. Tissue sample containing LV. C. Isolated and cleaned vessel for RNA extraction. D&E, Quantification of LVs inflammation was determined using the ratio of MPO staining to cell nuclei (DAPI), in this example 1.0%/6.3%= 0.16, to divide samples into low, medium and high inflammatory states with thresholds of <3%, 3-10% and >10% respectively.

### MiRNA extraction and analysis

Samples in RNAlater were individually defrosted and homogenized in 700μL of QIAzol (Qiagen, UK), using a PT 1300 D homogenisor with a 3mm dispersing aggregate head (Kinematica, Switzerland) at 20,000 RPM in a sterile 1.5 ml tube. RNA was extracted using a miRNeasy Mini Kit (Qiagen, UK) following the standard protocol and quality was verified using a NanoDropTM 2000 at A260/280nm and A260/230nm absorbance (Thermofisher, UK). Each sample was reverse-transcribed to cDNA using a miScript II RT Kit (Qiagen), then qPCR was carried out using a miScriptSybrGreen real-time PCR kit (Qiagen) and pre-defined miScript miRNA PCR Array (Qiagen, UK). The array contained primer pairs for 8 housekeeping genes and controls plus 88 human miRNAs that have previously been identified or predicted to target genes involved in the regulation of inflammatory responses and autoimmunity. One well was populated with a negative control sample. The array was run using a StepOnePlus Real-Time PCR System (Applied Biosystems), according to the manufacturer’s instructions.

### Immunofluorescence staining and imaging

Samples embedded in OCT were sectioned at 10-16μm and stored at -80°C. Both standard Haematoxylin and Eosin (H&E) and immunofluorescent staining were subsequently performed. For immunofluorescent staining, slides were washed with Tris Buffered Saline (TBS) (Sigma-Aldrich, UK) and 0.05% Triton-X (Sigma) and blocked for 1hour with 10% goat serum/TBS (Sigma-Aldrich). Slides were incubated overnight at 4°C with primary antibodies diluted in 5% goat serum/TBS at ratios of 1:200 for inflammatory marker Myeloperoxidase (MPO) (ab9535), 1:200 for ovarian carcinoma marker Wilms Tumour-1 (WT1) (ab89901), 1:50 for ovarian carcinoma marker Paired-Box-Gene 8 (Pax-8) (ab189249), and 1:200 for LEC marker Podoplanin (PDPN) (ab10288). Alexa-488 conjugated secondary antibody, anti-Mouse (ab150113) or anti-Rabbit (ab150077) were applied at a dilution of 1:250 in 5% goat serum/TBS at room temperature for 1 hour. Samples were stained with DAPI (4’,6-diamidino-2-phenylindole) solution (Sigma-Aldrich) and mounted with Fluoroshield (Sigma-Aldrich). Confocal Z-stacks and multi-channel images were processed and analysed using FIJI imaging software [24].

Assessment of H&E staining confirmed cryo-sections of LV samples stored at -80°C maintained sufficient morphology to quantify inflammation and cancer cell presence. The identification of LVs was confirmed by the positive staining with Podoplanin. The LV were subsequently classified into a low, medium or high inflammatory group with thresholds of <3%, 3-10% and >10% of MPO staining to nuclei (DAPI) staining ratio (Fig 1 D and E). The level of cancer-cell infiltration was determined by the number of cancer cells detected in a LV cross-section, as shown by positive staining with WT1 and Pax-8.

### Statistical analysis

Clinical correlation between different combinations of LV state (inflammation, high-inflammation, cancer-infiltration), and patient relapse, and cancer-stage were assessed by applying Fishers exact test, to identify significant non-random associations, and calculation of Cohen’s Kappa coefficient of inner agreement (κ), interpreted according to the methods of Vierra and Garett 2005 [25]. A P value below 0.05 was considered statistically significant. Kaplan Meier estimates of probability of non-relapse were also calculated for cancer-infiltration, inflammation and age. Prior to further analysis, the miRNA expression of each LV (ΔCT) was normalized to the expression of a stably expressed gene in each array, confirmed with NormFinder (http://moma.dk/normfinder-software) and GeNorm (http://www.biogazelle.com/qbaseplus) software. SNORD96 was selected as a reference “housekeeping” gene. The fold change between imaging-informed groups of LVs was calculated as ΔΔCT Group2/ΔΔCT Group1 where ΔΔCT=2^Δ^CT. Fold regulation was normalised to values greater than 1, noting up- or down-regulation. Principle component analysis (PCA) was used to transform multivariate data linearly into a set of uncorrelated variables to identify patterns in the variance amongst the data. We applied PCA to the miRNA expression recorded across all our samples to calculate two principle components (PC) that account for the greatest sources of miRNA variation within samples, using online software ClustVis (https://biit.cs.ut.ee/clustvis/) [26]. In cases where more than 1 miRNA was identified as significantly up-regulated or significantly down-regulated, pathway analysis was carried out using Diana-MiRPath v3 [27]. After removing miRNA that did not show a >1.8 fold-regulation change, a student-t-test, assuming equal variance, was applied to further confirm variation between groups. To visualise the distributions of the individual miRNA for which statistical significant difference was identified between groups, the probability density functions for miRNA expression in each group were estimated using Kernel Density Estimated distributions.

A number of widely used automatic classification methods were then used to predict the binary outcomes of the individuals (relapse vs. non-relapse, cancer infiltration vs. no cancer infiltration, medium or inflammation versus low inflammation), based on the LV miRNA expression of the individuals as predictor (input) variables. We used the supervised classification methods Logistic Regression, K-Nearest Neighbours, Support Vector Machine, Random Forest, and Gaussian Naïve Bayes. Evaluation of each method was performed using a leave-one-out cross validation procedure: each patient was considered in turn as a new patient (i.e. the test patient), while the remainder were considered as the training set. The predicted and actual outcome for each test patient was then compared with method accuracy measured as the percentage of correct predictions. Two other measures of accuracy were also used: the Cohen’s Kappa coefficient (κ) of inner agreement between predicted and actual values, and the McNemar’s test measuring the significance of equality of predicted probability (inner accuracy) between groups for each outcome. In each method, we also split the cases according to the binary outcome variable and measured the percentage of correct predictions in each group. A set of only the miRNA that were significantly differentially expressed between groups was then used to train new classifiers. We then assessed whether use of the new classifiers could improve predictions when compared to the performance of classifiers built using the entire miRNA expression set.

### Data availability

Further analytical results are available in the supplementary material. All raw and normalised expression data files have been deposited within Imperial College London’s Research Data Repository at http://doi.org/10.14469/hpc/7192.

## Results

### Post-operative assessment of patients

Ten patients were included in the present analysis. Nine of those patients had a high grade serous histology, and one patient had a clear cell carcinoma (Table 1). 60% of the lymph vessels were harvested at primary debulking surgery and 40% at interval debulking surgery (S1 Table). Only one positive para-aortic LN was detected at final histopathology. In the 13.2±3.6 months follow up period, 4 of the 10 patients had relapsed. Of the four patients that relapsed, three vessels demonstrated high inflammation and one vessel showed medium inflammation. Vessels from patients that did not relapse showed low inflammation in 4 cases and medium inflammation in the remaining 2 cases. Of the four relapse cases, cancer-cell infiltration was detected in three vessels (two with high inflammation and one showing medium inflammation). The overall relative range of inflammation in the lymphatic vessels was small with high inflammation defined as an average of >10% MPO:DAPI ratio and a narrow range of an average of 3-10% MPO:DAPI ratio for medium inflammation. All vessels showed less inflammation than similarly stained ovarian tumour tissue (Fig 2I). Consequently, to compare LVs with a notable difference in inflammation we focused initially on comparison of the LVs with high inflammation versus low inflammation.

**Fig 2.**
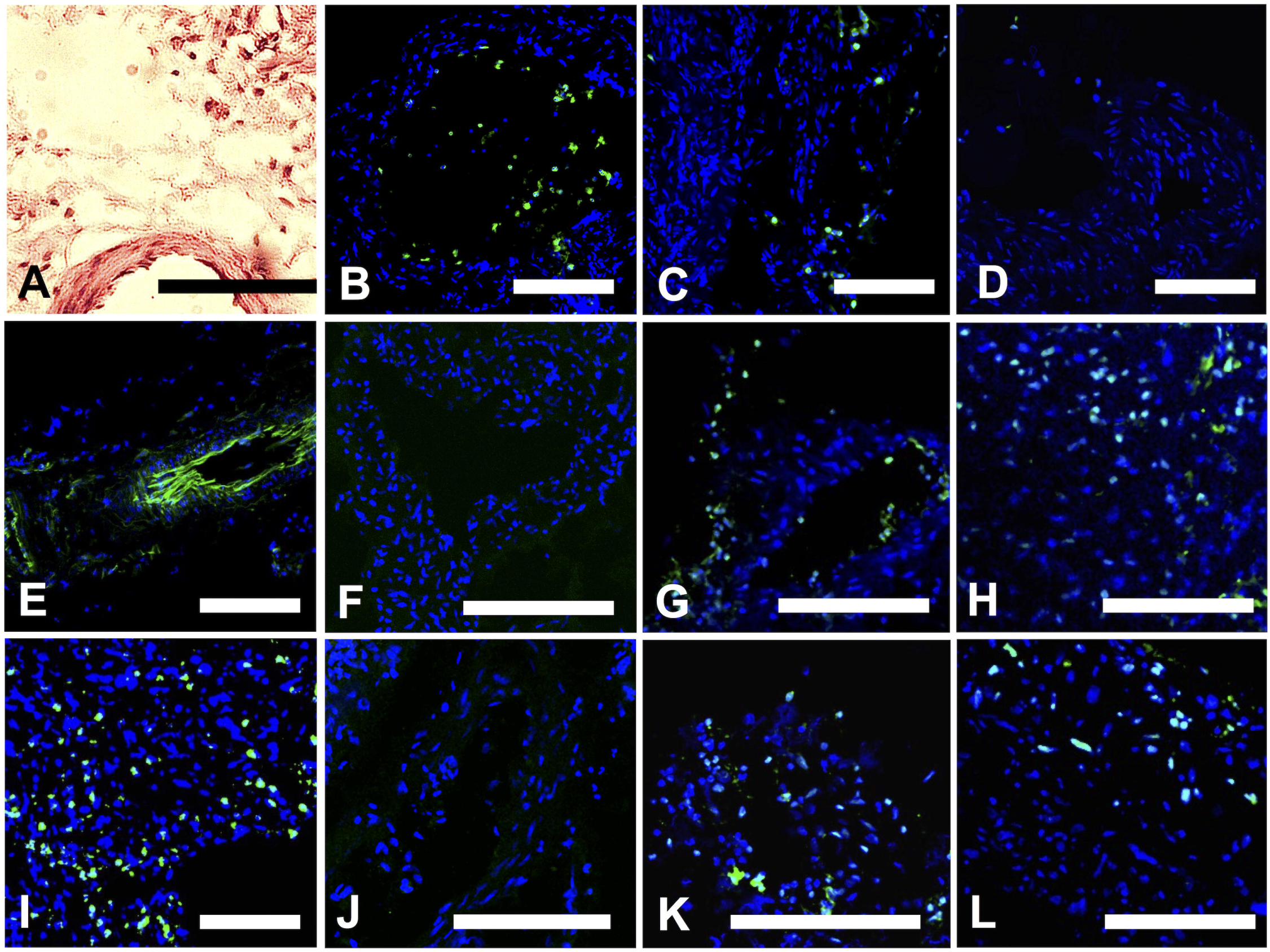
Lymphatic vessel imaging. A. H & E stained sections of the LVs, show that the nuclei were sufficiently preserved for subsequent Paired-Box-Gene 8 (Pax-8) and WT1 (Wilms-Tumour-1) staining. B-D. LVs with high, medium and low/no inflammatory state determined with myeloperoxidase (MPO) staining. E. Confirmation of LV status was carried out using LEC marker Podoplanin (PDPN) that stained around the LV lumen. F-H. Imaging with WT1, (F) negative LV, (G) positive LV (H) positive control. I. Tumour MPO positive control. J-L. Imaging with Pax-8, (J) negative LV, (K) positive LV and (L) positive control. An Ovarian tumour sample was used as a positive control and stained for both (M) WT1 and (Q) Pax-8. Scale=140μm.

### Clinical correlation of LV inflammatory state or cancer-cell infiltration and patient relapse

Of the four cancer-infiltrated LVs identified, 2 presented with high inflammation and the remaining 2 samples presented with medium inflammation. Cancer-free LVs presented with generally lower degrees of inflammation (Table 1). Tumour cells were detected in 50% of LVs with medium inflammation and two-thirds of LVs displaying high inflammation. The LVs presented with high inflammation in 3 of 10 samples, two of which were also positive for cancer cells. Three LV samples presented with medium inflammation, two of which contained cancer cells, and the remaining 4 LV samples presented with low inflammation and no detectable cancer cell infiltration (Table 1). The samples were macroscopically normal in appearance, however positive staining for the ovarian tumour markers WT1 and Pax8 was observed in four of the 10 samples collected (Fig 2 G,K). The LVs with cancer cells showed a moderate agreement (κ=0.583) with patient relapse but this was not significant (Table 2) (p=0.114). A moderate but statistically non-significant agreement was found between cancer-infiltrated LVs and Stage IV and inflammation (κ =0.583, 0.61 respectively).

**Table 2:** Cohen’s Kappa Coefficient for inner-agreement between different sample states (1=perfect agreement) and Fisher’s exact test for identification of significant non-random association (p<0.05) (n=10).

| **Cohen's Kappa Co-efficient** |  |  |  |  |  |
| --- | --- | --- | --- | --- | --- |
|  | Relapse* | Inflam. | High Inflam. | Cancer-cell inf. | Stage IV |
| Relapse | x | 0.615 | **0.783** | 0.583 | 0.583 |
| Inflam. | 0.615 | x | x | 0.615 | 0.615 |
| High Inflam. | **0.783** | x | x | 0.348 | 0.348 |
| Cancer-cell inf. | 0.583 | 0.615 | 0.348 | x | 0.583 |
| Stage IV | 0.583 | 0.615 | 0.348 | 0.583 | x |
| **Fishers Exact Test (p-value)** |  |  |  |  |  |
|  | Relapse* | Inflam. | High Inflam. | Cancer-cell inf. | Stage IV |
| Relapse | x | 0.071 | **0.033** | 0.114 | 0.114 |
| Inflam. | 0.071 | x | x | 0.071 | 0.071 |
| High Inflam. | **0.033** | x | x | 0.3 | 0.3 |
| Cancer-cell inf. | 0.114 | 0.071 | 0.3 | x | 0.114 |
| Stage IV | 0.114 | 0.071 | 0.3 | 0.114 | x |

Inflam = comparison of LVs with high or medium inflammation to low inflammation. High
Inflam. = comparison of LVs with high inflammation to low or medium inflammation.
Cancer-cell inf. = cancer-cell infiltration.
\* *Mean time since surgery= 13.2 months ± 3.6*

As the degree of inflammation in the LVs increased, so did correlation with patient relapse. The presence of LVs showing high inflammation significantly correlated with patient relapse (κ= 0.783, p=0.03) (Table 3). The presence of medium or high inflammation within LVs showed a moderate agreement with patient relapse (κ= 0.615, p=0.07) and patients with Stage IV, relapse and cancer-cell infiltration (κ=0.615, p=0.07) (Table 2). The presence of highly inflamed LVs compared to the presence of low inflammation only, resulted in a stronger correlation involving high inflammation status (Table 3). LVs exhibiting high degrees of inflammation showed near perfect agreement with patient relapse compared to those with low inflammatory state (n=7, κ=1.0, p = 0.029). Additionally, there were indications of a relationship between highly inflamed vessels with cancer-infiltration and Stage IV cancer (κ=0.695), but this lacked statistical significance (p=0.143). These results therefore support the earlier identified correlation between high inflammation and relapse. Kaplan Meier estimates of non-relapse according to inflammatory state, cancer-infiltration and age thresholds (>55, >60, >70, >75 years) indicate that probability of non-relapse is decreased in patients with LVs displaying medium or low inflammation and patients with cancer-infiltrated LVs (S2 Fig). Probability of non-relapse showed that patients >55 years age were more likely to relapse (S3 Fig).

**Table 3:** Comparison of LVs with high inflammation vs those with low inflammation only (n=7) using Cohen’s Kappa Coefficient and Fisher’s exact test, as described in Table 2.

|  | Relapse* | Cancer | Stage IV |
| --- | --- | --- | --- |
| Cohen's Kappa Co-efficient | **1** | 0.695 | 0.695 |
| Fishers Exact Test | **0.029** | 0.143 | 0.143 |

\* Mean time since surgery= 13.2 months $\pm 3.6$

### Lymphatic vessel miRNA expression during EOC

Expression analysis of a panel of miRNA involved in auto-immunity and inflammation was carried out. This showed that let-7 was differentially expressed in the highly-inflamed LVs (Table 4, S2 Table). Significant differential expression of 8 miRNA was identified when comparing miRNA expression of LVs presenting with high inflammation versus low inflammation (p<0.05, n=7). Of the 8 miRNA, 6 belonged to the let-7 family with miR-23a-3p and miR-23b-3p completing the set. Differential expression can also be clearly visualised when the estimated probability density functions, calculated using kernel density estimation for inner groups, for the differentially expressed individual miRNA are compared (S4 Fig). Correction for multiple testing by applying a post-hoc Bonferroni correction showed that differential expression of three miRNA, let-7b, let-7c and let-7d remained significant (p<0.0017). Comparison of miRNA expression in LVs displaying low inflammation versus medium or high inflammation showed a similar trend towards down-regulation of the let-7 family (Table 4, S3 Table). Six miRNA were down-regulated >1.8 fold with significant down-regulation of let-7g-5p and let-7i-5p (p<0.05).

**Table 4:**
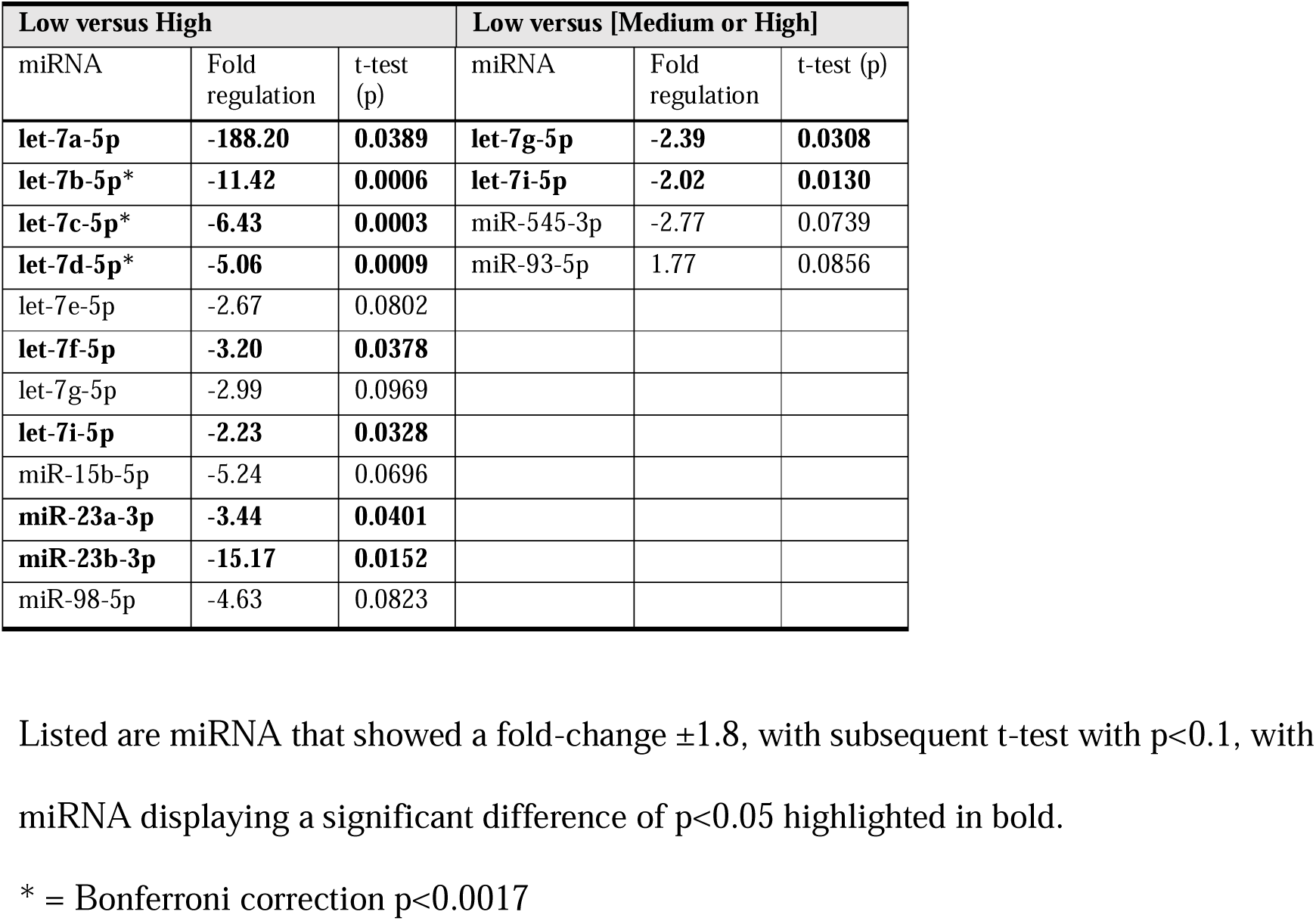
Differential expression of miRNA between LV groups displaying low, medium or high inflammation.

Similar trends were observed in the differential miRNA expression identified in the inflamed LVs and the differential miRNA expression in LVs collected from patients who underwent relapse during the study. Four significantly differentially expressed miRNA (p<0.05) were identified in relapse LVs (let-7c-5p, let-7b-5p, miR-23b-3p and miR-186-5p), with three of these (let-7c-5p, let-7b-5p, miR-23b-3p), also significantly differentially expressed in highly inflamed LVs (Table 5). Differential expression of miRNA with correction criteria of p<0.002 was not identified. Of the 17 miRNA that were down-regulated >1.8 fold in relapse vessels, 8 were also down-regulated in inflamed LVs, and 11 in the highly-inflamed LVs, including 5 and 6 members of the let-7 family respectively and miR-23b-3p (S2, S3 and S5 Table).

**Table 5:** Differential expression of miRNA between cancer-infiltrated versus non-cancer infiltrated LVs and LVs from relapse versus non-relapse patients within 13.2±6 months.

| Cancer-infiltrated LVs |  |  | LVs from patients that relapsed |  |  |
| --- | --- | --- | --- | --- | --- |
| miRNA | Fold regulation | t-test (p) | miRNA | Fold regulation | t-test (p) |
| **miR-144-3p\*** | **14.02** | **0.0010** | miR-144-3p | 5.07 | 0.0906 |
| **miR-181c-5p** | **10.63** | **0.0170** | **miR-186-5p** | **4.93** | **0.0294** |
| miR-101-3p | 5.08 | 0.0740 | miR-497-5p | 2.29 | 0.0672 |
| **miR-381-3p** | **4.07** | **0.0140** | miR-34a-5p | 2.09 | 0.0828 |
| **miR-497-5p** | **2.9** | **0.0260** | miR-98-5p | -3.43 | 0.0673 |
| **miR-93-5p\*** | **2.25** | **0.0010** | let-7d-5p | -3.68 | 0.0645 |
| **miR-16-5p** | **2.08** | **0.0310** | **let-7c-5p** | **-4.33** | **0.0385** |
| **let-7i-5p** | **-2.08** | **0.0210** | **let-7b-5p** | **-6.94** | **0.0383** |
| let-7b-5p | -2.27 | 0.0700 | **miR-23b-3p** | **-8.53** | **0.0278** |
| let-7c-5p | -2.56 | 0.0780 |  |  |  |

Listed are miRNA that showed a fold-change $\pm 1.8$ , with subsequent t-test with $p < 0.1$ , with miRNA displaying a significant difference of $p < 0.05$ highlighted in bold.
\* = Bonferroni correction $p < 0.002$

In the cancer-infiltrated LVs, most miRNA expression changes involved miRNA up-regulation such as miR-16-5p, miR-93-5p and miR-497-5p, miR-381-3p, miR-144-3p and miR-181a-5p (p<0.05) (Table 5). Of these, up-regulation of miR-93-5p and miR-144-3p remained significant with application of a Bonferroni correction (p<0.002). In contrast, inflamed LVs presented with primarily down-regulated miRNA, in particular, significant down-regulation of members of the let-7 family along with miR-23a-3p, miR-23b-3p and tumour suppressor miR-15-5p [28]. In the cancer-containing LVs, only 7 miRNA showed a >1.8 fold down-regulation, 6 of which were members of the let-7 family (S4 Table). Of these six, five were also down-regulated >1.8-fold in the LVs of patients who relapsed during the study period.

Similar trends in the differential miRNA expression in cancer-infiltrated LVs and highly inflamed LVs were also identified, as 15 miRNA were found to undergo an expression change of >1.8 fold (up- or down-regulated) in both of the comparisons (S2 and S4 Table). Six of the 7 miRNA that were down-regulated in both groups belonged to the let-7 family. This common trend in the fold-regulation difference in the highly inflamed and cancer-infiltrated LVs may be reflective of the fact that 50% of the cancer-infiltrated LVs also presented with high inflammation. We also noted that the number of differentially expressed miRNA was higher when comparing LVs presenting with high or low inflammation than when comparing vessels with or without cancer-infiltration. Expression of several miRNA were consistently below qPCR detection levels and thus omitted from further analysis.

### Pathway analysis of miRNA found to be dysregulated in inflamed or cancer-infiltrated LVs

Pathway analysis indicated that significantly dysregulated miRNAs are involved in maintaining LV integrity. The 7 miRNAs that were significantly upregulated in cancer-infiltrated LVs, and the 7 miRNAs that were significantly down-regulated in inflamed LVs have known involvement in various pathways related to the TGFβ-pathway, fatty acid biosynthesis, proteoglycans in cancer, glycosaminoglycan synthesis and glycan biosynthesis as well as the extracellular matrix receptor interactions pathway and adherens junction pathway (Table 6). No significantly up-regulated miRNAs were identified in highly inflamed LVs and only one significantly down-regulated miRNA was identified in cancer-infiltrated LVs. Pathway analysis carried out with the 4 significantly dysregulated miRNAs in the relapse LVs showed a range of potential target pathways including adherens junction, ECM-receptor interactions and mucin type O-Glycan biosynthesis (S6 Table).

**Table 6:** Pathway analysis using MiRNA that were significantly up-regulated in cancer-infiltrated LVs (left-hand side) and miRNA significantly down-regulated >1.8 fold (right-hand side) in highly inflamed LVs.

| Upregulated miRNA in cancer-infiltrated LVs |  |  |  | Down-regulated miRNA in highly inflamed LVs |  |  |  |
| --- | --- | --- | --- | --- | --- | --- | --- |
| KEGG pathway | p-value | Genes | miRNA | KEGG pathway | p-value | Genes | miRNA |
| TARBASE |  |  |  | TARBASE |  |  |  |
| Proteoglycans in cancer | 3.02E-15 | 96 | 5 | Adherens junction | 5.86E-13 | 52 | 8 |
| Adherens junction | 3.09E-09 | 44 | 5 | Proteoglycans in cancer | 5.86E-13 | 107 | 8 |
| Prion diseases | 3.09E-09 | 14 | 5 | TGF-β signaling pathway | 3.29E-09 | 49 | 8 |
| Viral carcinogenesis | 1.64E-08 | 91 | 5 | Viral carcinogenesis | 5.77E-09 | 99 | 8 |
| Hippo-signaling pathway | 2.37E-08 | 68 | 5 | Cell cycle | 2.79E-08 | 74 | 8 |
| TGF-β signaling pathway | 9.56E-08 | 41 | 5 | Hippo signaling pathway | 1.24E-07 | 74 | 8 |
| TARGETSCAN |  |  |  | TARGETSCAN |  |  |  |
| Fatty acid biosynthesis | 7.90E-38 | 2 | 2 | Fatty acid biosynthesis | 2.92E-53 | 3 | 1 |
| Fatty acid metabolism | 8.15E-15 | 3 | 2 | Fatty acid metabolism | 1.03E-21 | 4 | 1 |
| Metabolism of xenobiotics by cytochrome P450 | 7.48E-06 | 2 | 2 | Signaling pathways regulating pluri-potency of stem cells | 2.58E-09 | 17 | 7 |
| Signaling pathways regulating pluri-potency of stem cells | 6.26E-05 | 11 | 3 | Metabolism of xenobiotics by cytochrome P450 | 8.99E-07 | 3 | 3 |
| TGF-β signaling pathway | 0.0387 | 5 | 3 | N-Glycan biosynthesis | 0.00466 | 5 | 3 |
| Micro-CT-DS |  |  |  | Micro-CT-DS |  |  |  |
| Fatty acid biosynthesis | 7.90E-38 | 5 | 2 | Fatty acid biosynthesis | 7.39E-26 | 6 | 9 |
| Fatty acid metabolism | 8.15E-15 | 11 | 2 | ECM-receptor interaction | 3.21E-07 | 17 | 9 |
| Signaling pathways regulating pluri-potency of stem cells | 3.74E-12 | 47 | 3 | Signaling pathways regulating pluri-potency of stem cells | 1.01E-06 | 43 | 9 |
| Hippo signaling pathway | 2.06E-06 | 29 | 2 | Mucin type O-Glycan biosynthesis | 5.38E-06 | 9 | 9 |
| Proteoglycans in cancer | 2.70E-06 | 59 | 4 | Proteoglycans in cancer | 2.34E-05 | 53 | 9 |
|  |  |  |  | p53 signaling pathway | 3.13E-05 | 27 | 9 |

### Principle component analysis of miRNA expression in LV samples

PCA showed that differential miRNA expression identified in inflamed and cancer-infiltrated LVs could also be identified without grouping the samples. This can be seen when assessing the top 16 miRNA whose expression contributed the most to the first and second Principle Components (PC1 and PC2). These components capture factors responsible for the most and second most variation in the data set, respectively. Samples with the most similar miRNA expression profile cluster together and clusters may vary over PC1/PC2 or both (S5 Fig). Twelve miRNA identified in the fold-change analysis of highly inflamed or cancer-infiltrated LVs, were among the top 20 miRNA that contributed to PC1, with 5 of those showing significant differential expression. Furthermore, 8 miRNAs of the top 20 miRNA that contributed to PC2 were all identified as significantly downregulated >1.8 fold in highly inflamed LVs. The remaining miRNA varied greatly between the patient samples, e.g., miR-20, but were not identified by analysis of fold change between groups. The expression levels of these miRNA were therefore spread equally between the compared groups. Overall, variation over PC2 differentiates between highly-inflamed and non-inflamed LVs, supporting the suggestion that miRNA expression could distinguish LV states (S5 Fig).

### Application of machine-learning algorithms to predict LV state or clinical outcome based on miRNA expression

Patterns of miRNA expression showed some potential in predicting clinical outcomes by applying classification algorithms. When predicting patient relapse (Table 7), all of the classification methods presented with quite high accuracy values and McNemar’s probabilities of p<0.05, meaning that no significant difference was found between the predicted grouping, based on miRNA expression, and the grouping based on the known clinical data. Of the methods applied, use of k-Nearest Neighbours and logistic regression produced the highest classification accuracy with similar prediction accuracy achieved between methods. Classification of cancer-cell infiltrated LVs based on miRNA expression (Table 8) showed a similar prediction accuracy compared to classification of relapse LVs, but the Random Forest classifier achieved the best accuracy. Classification of inflamed and <=Stage IV samples did not show a high accuracy (S7 Table). As a further evaluation, we examined the effect of keeping as predictors only those miRNAs that have significant associations with the outcome variables. Analysis was performed on classifiers build using only significantly differentially expressed miRNAs (p<0.05). In the cancer-infiltrated LVs, the differentially expressed miRNAs enhanced the accuracy of the predictive algorithm to 80-100% (S8 Table). Prediction of relapse based on miR-23b-3p and miR-186-5p expression, improved the accuracy of classification with all methods by 10%, except when using the Support Vector Machine, where no improvement was observed (see S8 Table & S1 File). The addition of miR-let-7b-5p and let-7c-5p improved the accuracy of the Support Vector Machine from 80% to 90% but decreased the accuracy of Logistic Regression from 80% to 70%. However, the use of miR-23b and miR-186-5p only showed a bias towards classifying the samples as non-relapsed when using Logistic Regression, which decreased when using all 4 miRNA. These results suggest that the miRNAs with significant associations to the outcome variables have a capacity to be used as predictors for predicting the outcomes of interest in new patients.

**Table 7:** Classification analysis performance for various supervised classification algorithms for the prediction of patient relapse based on LV miRNA expression. This illustrates the accuracy of the method (percentage of correct predictions), the per-group accuracy (No Relapse, Yes Relapse), Cohen’s Kappa of inner agreement between predicted and actual outcome, and McNemar’s test of significance of equality of predicted probability (inner accuracy) between groups for each outcome.

| Method | Accuracy | McNemar's Test-pval | Cohen's Kappa | No Relapse | Yes Relapse |
| --- | --- | --- | --- | --- | --- |
| Logistic Regression | 70% | 0.999 | 0.375 | 67 % | 75 % |
| K-Nearest Neighbours | 80% | 0.289 | 0.583 | 100 % | 50 % |
| Support Vector Machines | 80% | 0.289 | 0.583 | 100 % | 50 % |
| Random Forests Classifier | 50% | 0.375 | -0.042 | 67 % | 25 % |
| Gaussian Naive Bayes | 60% | 0.031 | 0.167 | 100 % | 0 % |

**Table 8:** Classification performance of various supervised classification algorithms for the prediction of cancer infiltration based on LV miRNA expression. Details are similar to Table 7.

| Method | Accuracy | McNemar's Test-pval | Cohen's Kappa | No Cancer | Yes Cancer |
| --- | --- | --- | --- | --- | --- |
| Logistic Regression | 70% | 0.44 | 0.38 | 83 % | 50 % |
| K-Nearest Neighbours | 50% | 0.37 | -0.04 | 67 % | 25 % |
| Support Vector Machines | 70% | 0.13 | 0.37 | 100 % | 25 % |
| Random Forests Classifier | 100% | 1.00 | 1.00 | 100 % | 100 % |
| Gaussian Naive Bayes | 50% | 0.37 | -0.04 | 67 % | 25 % |

## Discussion

To our knowledge, this is the first analysis of inflammatory and miRNA alterations in tumour draining LVs in ovarian cancer. In this feasibility study, we aimed to provide insights into LV features that might be associated with patient relapse. Our results may contribute to a better prediction of patient relapse, and could guide future experimental work seeking to improve patient outcome via therapeutic targeting of miRNA.

Macroscopically normal LVs, remote from the tumour site that exhibited high or medium inflammation correlated with cancer relapse. Despite cancer-cell infiltration in 66% of these vessels, a moderate but non-significant association was identified between inflamed and cancer-infiltrated LVs (p=0.07). The staging of LNs is routinely used to quantify cancer progression but the status of the connecting LVs has not previously been investigated. While case numbers are small, the presence of high inflammation in the LVs showed the strongest significant correlation with cancer relapse (p=0.03 κ=0.783). The Kaplan Meier estimates of non-relapse also showed that relapse became less likely without the presence of LV inflammation or cancer-cell infiltration, and age did not increase likelihood of relapse (S Fig 2 and 3). Whether inflammation is a consequence of factors that contribute to relapse, or the inflammation-induced changes directly affect relapse, is an area for future work.

Analysis of the effect of cancer cell infiltration of the LVs, without considering LV inflammatory state, identified a trend towards a non-significant association with relapse (p=0.07, κ=0.783). This suggests that cancer-infiltration of LVs alone may not necessarily predict relapse, but rather accompanying changes in vessel inflammation may increase the likelihood of relapse. Furthermore, cancer-cell infiltration did not show a significant association with high inflammation. The lack of a significant correlation in both cases may also be simply due to the possible scenario that cancer cells are just transiting through these vessels, so detecting their presence becomes a matter of timing. Overall, this study suggests that nearby LVs appear to play an important role in the immune response to cancer and the association between inflammatory lymphatic changes and relapse deserves further exploration while potentially representing a therapeutic target.

The use of miRNA expression classification algorithms also showed potential to predict relapse, with relatively high accuracy for these 10 patients when applying a k-Nearest Neighbours algorithm or a Support Vector Machine model (80%), rising to 90% when using significantly differentially expressed miRNA only (Table 7, S8). Further studies spanning a larger number of patients should therefore be performed. Clinically, the prediction of a patient’s outcome based on miRNA expression of tissue collected during surgery could be of considerable value. Despite the presence of inflammation showing a moderate agreement with relapse and high inflammation displaying a significant correlation with relapse, a classifier based on miRNA expression in high or medium inflamed LVs did not show a high accuracy (Tables 2 and 3, S7 and S8 Table). Furthermore, due to the limited number of highly inflamed LVs, it was not possible to build a reliable classifier based on miRNA expression in LVs with high inflammation versus miRNA expression in LVs with low or medium inflammation. Indeed, all the classifiers are limited by the number of samples required to train the predictive algorithms. Therefore, whilst providing insight, these observations require increased sample input to maintain confidence in miRNA classification.

Analysis of miRNA expression of highly inflamed LVs compared to LVs with no or low inflammation, showed down-regulation of 6 members of the let-7 family, 3 of which remained significant with a Bonferroni correction (p<0.0017). A similar trend was observed when comparing miRNA expression of LVs presenting with high or medium inflammation to those with low inflammation. In this case 6 members of the let-7 family showed down-regulation >1.8 fold but only two members showed significant down-regulation (p<0.05), which may be reflective of less difference in mean LV inflammatory state between the two groups compared. The change in let-7 miRNA expression was large enough that without designating groups, the overall variability in sample miRNA expression due to let-7 expression could be detected using PCA (S5 Fig). In ovarian tumour tissue, let-7 expression is down-regulated and associated with poor prognosis and chemo-resistance [17,29,30]. Let-7 family members target oncogenes HMAG2 and RAS, and loss of let-7 is also associated with the development of aggressive cancers [31]. Chemo-sensitivity of tumours may also be affected by let-7 expression. Manipulation of let-7e in EOC cells *in-vitro* and *in-vivo* confirmed that expression is reduced in cisplatin-resistant cell lines and application of let-7e mimics alongside cisplatin, reduced tumour growth in mice more than with cisplatin alone [32]. Furthermore, chimeric-mediated delivery of let-7i miRNA to EOC cells *in-vitro* reversed paclitaxel-induced chemo-resistance [33]. Expression of let-7 is also associated with control of the Cdk-dependent cell cycle network, that drives the cell transformation network, allowing cells to undergo an epigenetic switch between non-transformed and a malignant transformed state [34].

The implications of let-7 down-regulation in LVs include changes in immune response and LV permeability. Members of the let-7 family can target inflammatory cytokines, such as IL-6 by let-7a and IL-13 by let-7f, shown to contribute to cancer-cell apoptosis and anti-inflammatory action respectively [35,36]. In addition to affecting immune cells in transit, drainage of cytokines to LNs could impact immune response at the LN, potentially contributing to the loss of immuno-suppression observed in draining LNs [37]. Cytokines may also affect LV transport, as IL-6 can increase LV permeability while intradermal administration of cytokines IL-1β, TNF-α, and IL-6 have been shown to decrease lymphatic contraction frequency [38,39]. Down-regulation of let-7 in vascular endothelial cells is associated with increased TGF-β signalling leading to promotion of epithelial-to-mesenchymal transition and thus increased vessel permeability, but the effects in LVs are currently unknown [40]. The observed up-regulation of several miRNA (miR-381-3p, miR-93-5p, miR-16-5p and miR-497-5p) in cancer-infiltrated LVs may inhibit Vascular Endothelial Growth Factor (VEGF) expression or VEGF receptors, both directly and indirectly via Bcl-2 inhibition [41-44]. The inhibition of VEGF secretion in tumours has been a target of several drug trials but manipulation of VEGF secretion in connecting LVs may present as a future therapeutic area [45]. Expression of miR-144 has been shown to target RUNX-1 in EOC lines and SRF in HUVECs, both contributing to anti-proliferative action. A more extensive list of potential targets is provided in Table 9. A logical progression of this study would be to confirm targets in LECs.

**Table 9:**
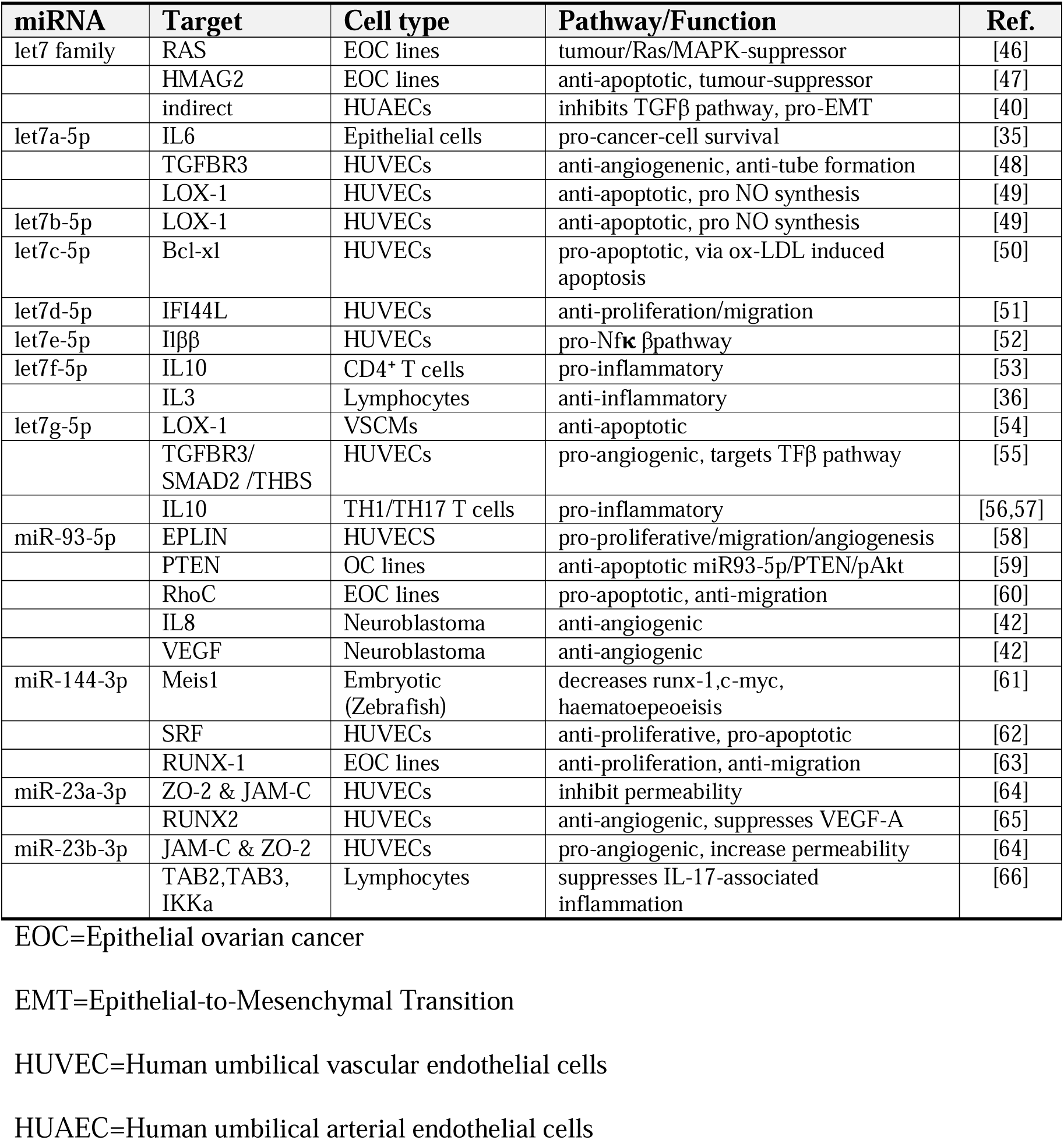
Confirmed targets of miRNA in primarily endothelial and cancer cell lines identified as significantly differentially regulated between groups.

Further studies of the relationship between inflammatory markers in LVs and differential miRNA expression would increase the potential for diagnostic capabilities. Jones et al. and Yee et al, analyzed LEC responses to inflammation and identified a total of 16 differentially expressed miRNA, one of which (miR-17-5p) overlapped with the panel of miRNAs screened in our study [18,19]. Chakraborty et al, identified differentially expressed miRNA associated with autoimmunity and inflammation in rat LECs, with different sets of miRNAs identified at 2 hours, 48 hours and 96 hours after induction of inflammation [13]. Our results showed no overlap with the miRNAs regulated at the 2-hour time-point but did show similar upregulation of miR-19a detected at 48 hours and induction of miR-497, miR-19b and miR-19a observed at 96 hours [13]. Additionally, there was no evidence of dysregulation of the let-7 family in the rat LECs. Taken together, chronic inflammation of LVs in EOC patients may impact miRNA expression differently, aligning more closely with the later time-points post-inflammatory induction in the controlled rat experiments.

The pathway analysis of the dysregulated miRNA identified involvement in transforming-growth factor TGF-β2 signalling, as well as glycoproteins in cancer and molecules that contribute to the glycocalyx of the LV lumen (Table 6). LECs possess a glycocalyx composed of glycoprotein side chains, a backbone that includes adhesion molecules, and a proteoglycan core. Therefore, changes to glycoproteins or adherens junctions may affect integrity of the LV wall [67]. Furthermore, a study by Zolla *et al* showed aged collecting LVs in rats presented with a thinner endothelial cell glycocalyx and a loss of gap junction proteins [68]. Functionally this resulted in hyper-permeability of the LVs that allowed pathogens to escape into surrounding tissue. This may suggest that the miRNA changes that accompany cancer cell infiltration and inflammation may also alter LV permeability and barrier function. This in turn could aid cancer cell infiltration but may also impact similar movements of immune cells such as macrophages and dendritic cells [69]. Further studies to assess the impact of inflammation-induced miRNA expression changes on LV integrity may identify new avenues to prevent transmigration of cancer cells across LV walls.

The response to cancer may also be mediated by inflammatory cells normally present in the LV. Analysis of rat mesenteric, pre-nodal and post-nodal LVs has shown the presence of innate inflammatory immune cells accumulating on the walls of the LVs including neutrophils, monocytes and macrophages [69]. Microarray analysis of mesenteric collecting vessels has also shown significant accumulation of MHCII+ antigen-presenting cells [70]. Furthermore, both antigen-presenting cells and T cells are transported in LVs, with proportions depending on the status of the upstream tissues [71]. We have not assessed the status of all of the tissues drained by these LVs, but in any case we would expect some immune cells to be captured in our images. However, the overall proportion of T cells to LV wall structural cells should be low enough not to impact overall miRNA expression.

The expression of the enzyme myeloperoxidase (MPO) was selected as an inflammatory marker due to evidence supporting MPO as a reliable inflammatory marker in a range of pathogenesis including cancer, rheumatoid arthritis and cardiovascular disease in addition to tissue injury [72]. The expression of MPO has also been associated with ovarian epithelial carcinoma cells in early stage carcinomas [73]. Presence of MPO is also specifically associated with vascular vessel inflammation [74,75]. Furthermore, MPO has been used as a reliable indicator of lymphatic vessel inflammatory status to quantify LPS-induced lymphatic vessel inflammation in rats in conjunction with Masson Trichome staining to indicate fibrotic vessel areas [13]. Other indicators of inflammatory state could be used, perhaps benefitting the prognostic value for relapse.

A significant limitation of our feasibility study is the small sample size. Although we have successfully identified associations with high potential for outcomes prediction, further samples could both strengthen the trends observed and uncover more obscure alterations associated with inflammation and/or relapse. Additionally, it is possible that increasing sample size could result in the identification of inflamed but non-cancer infiltrated LVs. However, there is no guarantee that increasing sample size would lead to the emergence of this group. The sample size also contributed to the decision to perform real time PCR with a relatively small number of pre-selected targets. A broader microarray study would require many more samples to avoid false positives. We screened for miRNA with known involvement in inflammatory and autoimmunity due to the previous characterization of inflammatory vessel state and pathophysiological changes associated with inflammation and lymphatic function in rat lymphatic vessels [13].

A further limitation to this study is the lack of samples from healthy controls. Harvesting equivalent lymphatic tissues from healthy patients would not be ethical. However, this means that the changes in LV status noted here are more directly relevant to identification of factors that aid prognosis than identifying systemic EOC changes compared to healthy subjects. Epithelial ovarian cancer surgical techniques and types of cytotoxic treatments do not differ, as of yet, between the various histological subtypes. We therefore included all available epithelial subtypes, even the clear cell case, to have a broader representation of the clinical reality. Although our results support a link between systemic inflammation and cancer cell dissemination, a larger study is warranted to validate our findings and to allow further analysis to better identify miRNA patterns using predictive algorithms with greater confidence.

## Conclusions

In conclusion, we have demonstrated that LVs appear to show quantifiable inflammatory and infiltrative alterations in patients with advanced EOC and that a high inflammatory LV state significantly correlates with higher risk of post-surgical relapse. Macroscopically normal lymphatic LVs show accompanying alterations in miRNA expression that may impact LV integrity and secretion of VEGF and inflammatory cytokines. Whether LV inflammation is a cause or consequence of cancer-cell infiltration is yet to be determined, but our results suggest systemic inflammation as a new area to target.

## Supporting information

**S1 File. Supplementary information regarding building supervised classification algorithms**. The techniques and assumptions when building classification algorithms and assessing the impact of individual miRNA.

**S1 Table. Additional clinical data (patient age and surgery type)**.

**S1 Fig. Experimental design for LV miRNA analysis from sample collection to analysis**. Each vessel dissected from one sample is divided into 2 and processed for IHC and miRNA expression analysis.

**S2 Fig. Kaplan Meier estimates of non-relapse (survival). A. According to inflammation (MPO) (high vs. medium/low) B. According to inflammation (MPO) (high/medium vs. low). C. According to cancer cell infiltration (WT1 or Pax**.**8**)

**S3 Fig. Kaplan Meier estimates of non-relapse (survival) according to multiple patient age thresholds. A. Under 55 years B. Under 60 years C. Under 65 years**. **D Under 70years. E Under 75years**.

**S2 Table. Differential expression of miRNA between LVs displaying high or low inflammation**. We compared expression in LVs with high versus low inflammation (n=7) Listed are miRNA that showed a fold-regulation change ±1.8 with those showing a significant difference between groups highlighted (t-test p>0.05). * = miRNA that remained below a Bonferroni correction of p<0.00208.

**S3 Table. Differential expression of miRNA between LVs displaying high or low inflammation**. We compared expression in LVs with high or medium inflammation versus low inflammation (n=10) Listed are miRNA that showed a fold-regulation change ±1.8 with those showing a significant difference between groups highlighted (t-test p>0.05). * = miRNA that remained below a Bonferroni correction of p<0.00161.

**S4 Table. Differential expression of miRNA between cancer-infiltrated LVs and non-cancer-infiltrated LVs**. Listed are miRNA that showed a fold-regulation change ±1.8 with those showing a significant difference between groups highlighted (t-test p>0.05). * = miRNA that remained below a Bonferroni correction of p<0.002.

**S5 Table. Differential expression of miRNA in LVs from patients that relapsed versus LVs from patients that didn’t relapse within 13**.**2**±**6 months**. Listed are miRNA that showed a fold-regulation change ±1.8 with those showing a significant difference between groups highlighted (t-test p>0.05). Bonferroni correction of p<0.002.

**S4 Fig. The KDE estimations of the distributions of inner groups for the individual miRNA expressed at statistically significant levels (p<0**.**05) between LVs with high inflammation and LVs with low inflammation**.

**S5 Fig. A. Principle component analysis showed that inflamed and non-inflamed samples show variance across Principle Component (PC) 2**. Samples with most similar miRNA expression profile cluster together. Unit variance scaling was applied to rows; SVD with imputation was used to calculate principal components. Prediction ellipses are such that with probability 0.95, a new observation from the same group will fall inside the ellipse. N = 10 data points. Figures produced with ClustVis [26] B. The miRNA expression driving PC1 & 2 contributed to the greatest variation between sample groups with several miRNA identified as differentially regulated when comparing inflamed or cancer-infiltrated LVs (bold =>±1.8, *=p<0.05). C. A Scree plot showing the amount of variation described by each PC confirmed that the two primary PCs accounted for much of the variation between all sample.

**S6 Table. Table of classification analysis for various supervised classification algorithms to predict LV inflammation (low or medium/high) or patient stage from LV miRNA expression**. Depicted is the accuracy of the algorithm to predict LV inflammation or patient relapse (percentage of correct predictions), the per-group accuracy (No Inflam., Yes Inflam. / Stage < IV, Stage >= IV), Cohen’s Kappa and thus the agreement between predicted and actual states, and McNemar’s test significance of equality of predicted probability (inner accuracy) between groups for each outcome.

**S7 Table. Classifier predictions of the examined states (LV inflammation, LV cancer-cell infiltration, patient relapse or patient stage) based on the expression of significantly dysregulated miRNA**. Accuracy, Cohen’s Kappa, Mc Nemar p-value of equality for inner grouped probabilities and classification are reported for the predicted classes. The expression of the most significantly differentially expressed miRNA were added one by one, or in pairs if significance was equal, into the parameter set used to build the classifiers. When only significantly differentially expressed miRNA identified in inflamed LVs (high and medium inflammation versus low) were used to build the classifier, the accuracy of subsequent predictions increased by 20-40% compared to classifiers based on all available miRNA expression (S6 Table). Similar improvements were found in the accuracy of classifiers predicting relapse and LV cancer-infiltration (Tables 7 and 8).

**S8 Table: Pathway-analysis performed with the 3 significantly up-regulated and single significantly down-regulated miRNA identified in the relapse versus non-relapse groupings**.

## Acknowledgements

The authors gratefully acknowledge the provision of tissue samples by the Imperial College Healthcare NHS Trust Tissue Bank. Other investigators may have received samples from these same tissues. The research was supported by the Experimental Cancer Medicine Centre (ECMC, Hammersmith Campus) and National Institute for Health Research (NIHR) Biomedical Research Centre based at Imperial College Healthcare NHS Trust and Imperial College London. The views expressed are those of the author(s) and not necessarily those of the NHS, the NIHR or the Department of Health.

