## Supplementary File 1 for "Inflammatory state of lymphatic vessels and miRNA profiles associated with relapse in ovarian cancer patients"

Supplementary Text.

### Building classification algorithms

Model parameters were estimated for k-nearest neighbours (k-NN), Support Vector Machines (SVM) and Random Forest Classifier using a grid optimization algorithm. For k-NN, the number of nearest neighbours was set to k=3 and for the SVM, a kernel type equal to 'rbf', Kernal gamma set to 0.0001, C value of 100 and Convergence epsilon of 0.001 was used. For the Random Forest classifier, number of estimators was equal to 7 with an entropy criterion. For Logistic Regression, the Wald Chi-square test of significance of estimators was > 0.05, indicating that the estimators are forming a statistical significant model for classification.

### Assessment of indidual miRNA

Two statistically significant individual miRNAs (let-7i-5p and let-7g-5p) were identified as differentially expressed between LVs with low inflammation and LVs with medium/high inflammation. When the classifier was built based on only let-7i-5p expression instead of the expression of all miRNA, accuracy was not increased. Use of Logistic Regression and SVM produced the lowest accuracy (60%) while k-NN and Gaussian Naive Bayes showed the highest (80%). Insertion of let-7g-5p increased accuracy using Logistic Regression and SVM (up to 80%) while insertion with use of the other methods resulted in no change in accuracy. For the 1 individual miRNA, Gaussian Naive Bayes and k-NN appear to be the least biased classification algorithms while use of Logistic Regression and SVM overestimated the samples as Inflamed. The same observation was noted when using both let-7i-5p and let-7g- 5p, where the algorithms tended to classify the samples as Inflamed, except when using the Gaussian Naive Bayes method, which showed a balance in predicted outputs.

The classifiers trained to predict cancer-infiltration based on differential miRNA expression presented with the highest accuracy of all classifiers tested. Seven individual miRNA showed significantly different distributions in cancer-infiltrated LVs (hsa-miR-144-3p, hsa-miR-93- 5p, hsa-miR-381-3p, hsa-miR-181-5c, hsa-let-7i-5p, hsa-miR-497-5p and hsa-miR-16-5p). The initial 2 miRNA presented with similar p-values, as did the 4th and 5th, therefore the miRNA expressions were added to the classifier in pairs. The classification algorithms such as Random Forest Classifiers and Gaussian Naive Bayes produced results reaching 100% accuracy of predictions, for all combinations of individual miRNAs, giving evidence that under a bigger sample, these classifiers can be trained efficiently and give high accuracy metrics. All the results showed an increasing rate of accuracy metric as miRNA were added into the analysis. When using all statistically significant miRNA four out of five classifiers reached 100% accuracy in prediction. Logistic Regression showed the lowest accuracy and appeared to be biased in classifying samples as non-cancer infiltrated. The SVM classifier also seemed to classify the samples as non-cancer infiltrated when using at least 5 examined individual miRNAs.
