## Supplementary Table 1 for "Inflammatory state of lymphatic vessels and miRNA profiles associated with relapse in ovarian cancer patients"

| Sample | Age | Surgery | Residual Disease | LN sampled | LN histology |
|---|---|---|---|---|---|
| 1 | 54 | Primary | nil | Right pelvic | negative |
| 2 | 57 | Primary | nil | No | n/a |
| 3 | 62 | Interval | nil | No | n/a |
| 4 | 64 | Interval | nil | No | n/a |
| 5 | 57 | Primary | nil | No | n/a |
| 6 | 61 | Interval | nil | No | n/a |
| 7 | 81 | primary | nil | Para-aortical | positive |
| 8 | 41 | Interval | nil | No | n/a |
| 9 | 75 | Primary | nil | No | n/a |
| 10 | 64 | primary | nil | Left pelvic &para-aortic | both negative |
