## Supplementary figures and images for "Inflammatory state of lymphatic vessels and miRNA profiles associated with relapse in ovarian cancer patients"

### Supplementary Figure 1

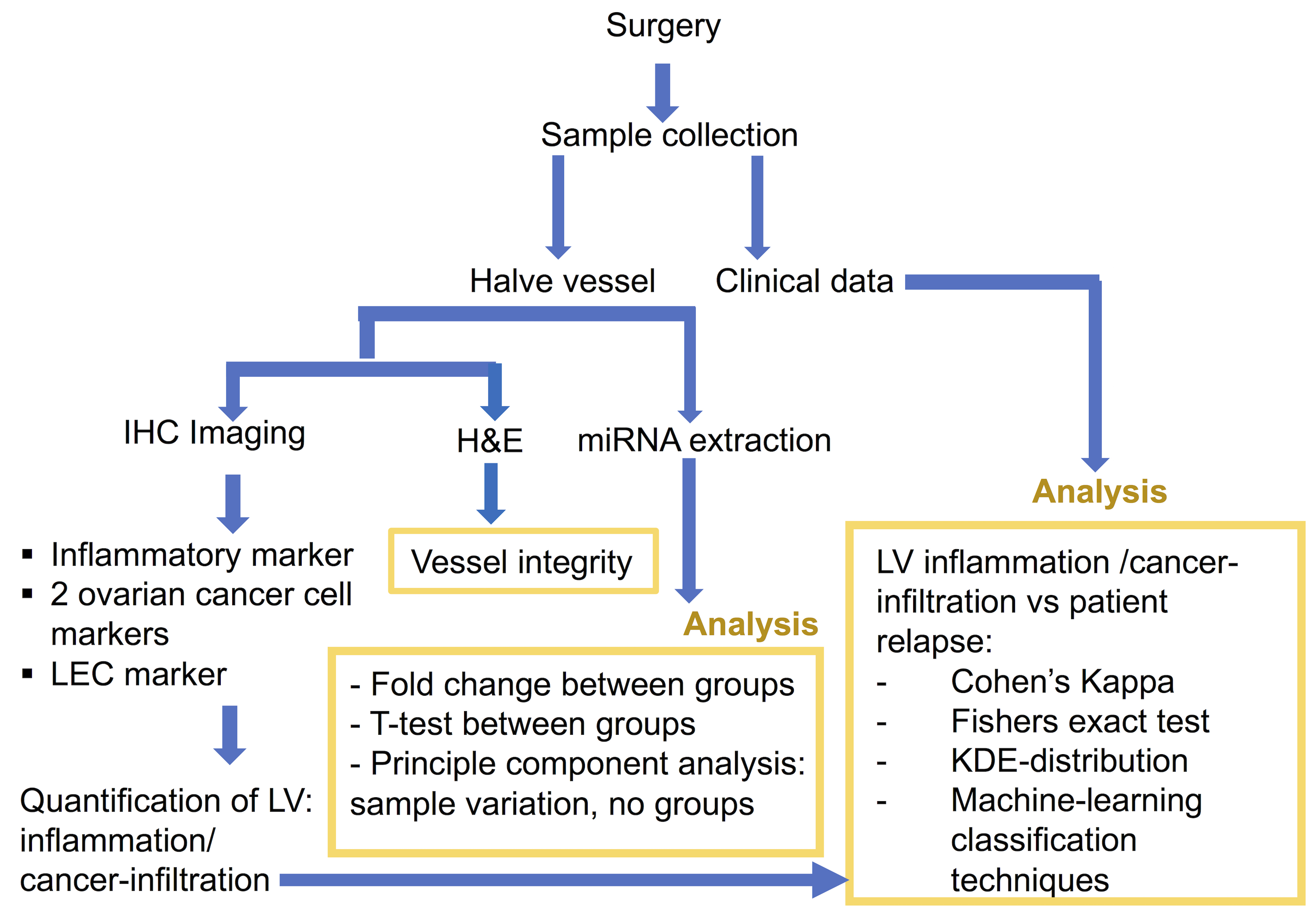

### Supplementary Figure 2

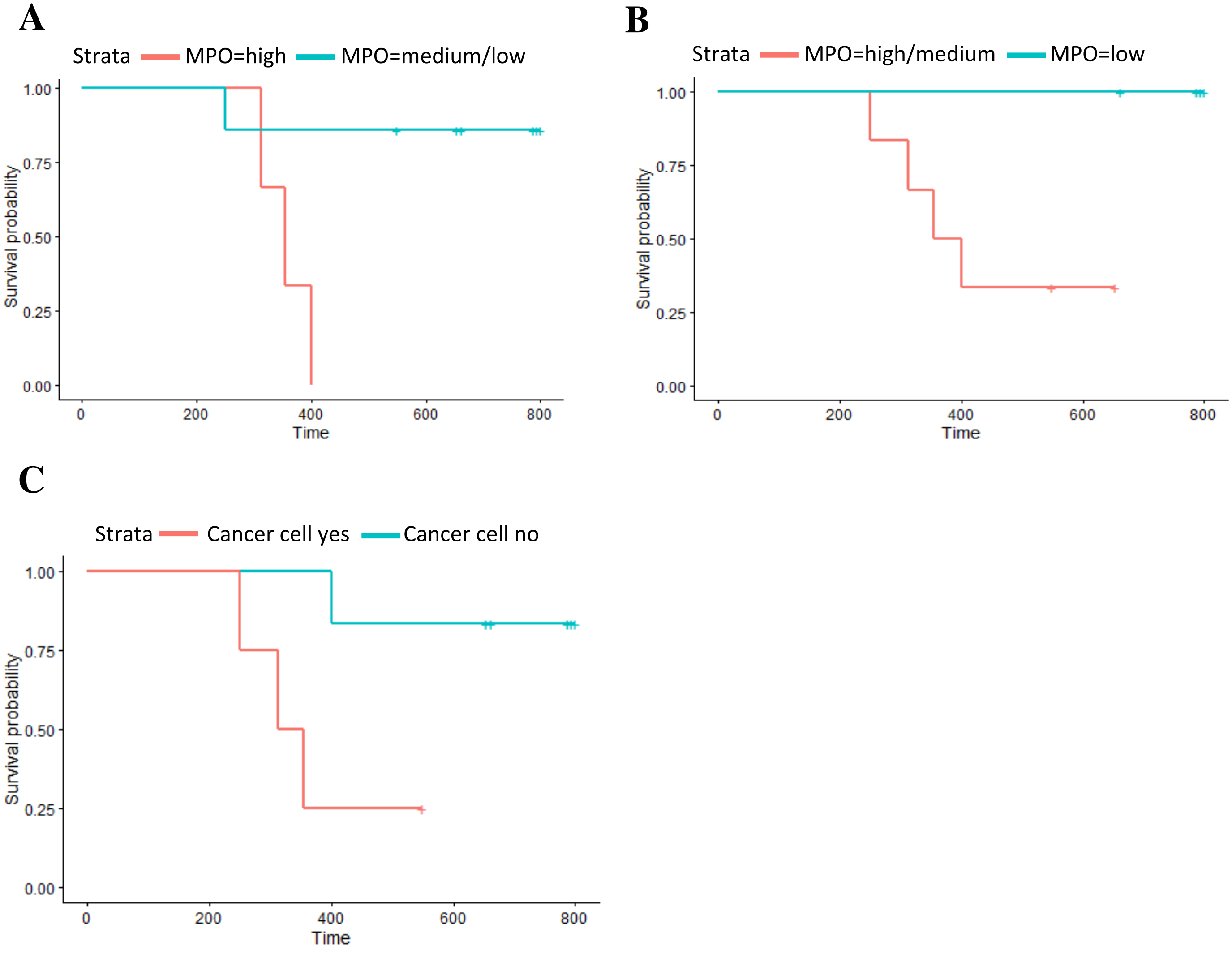

### Supplementary Figure 3

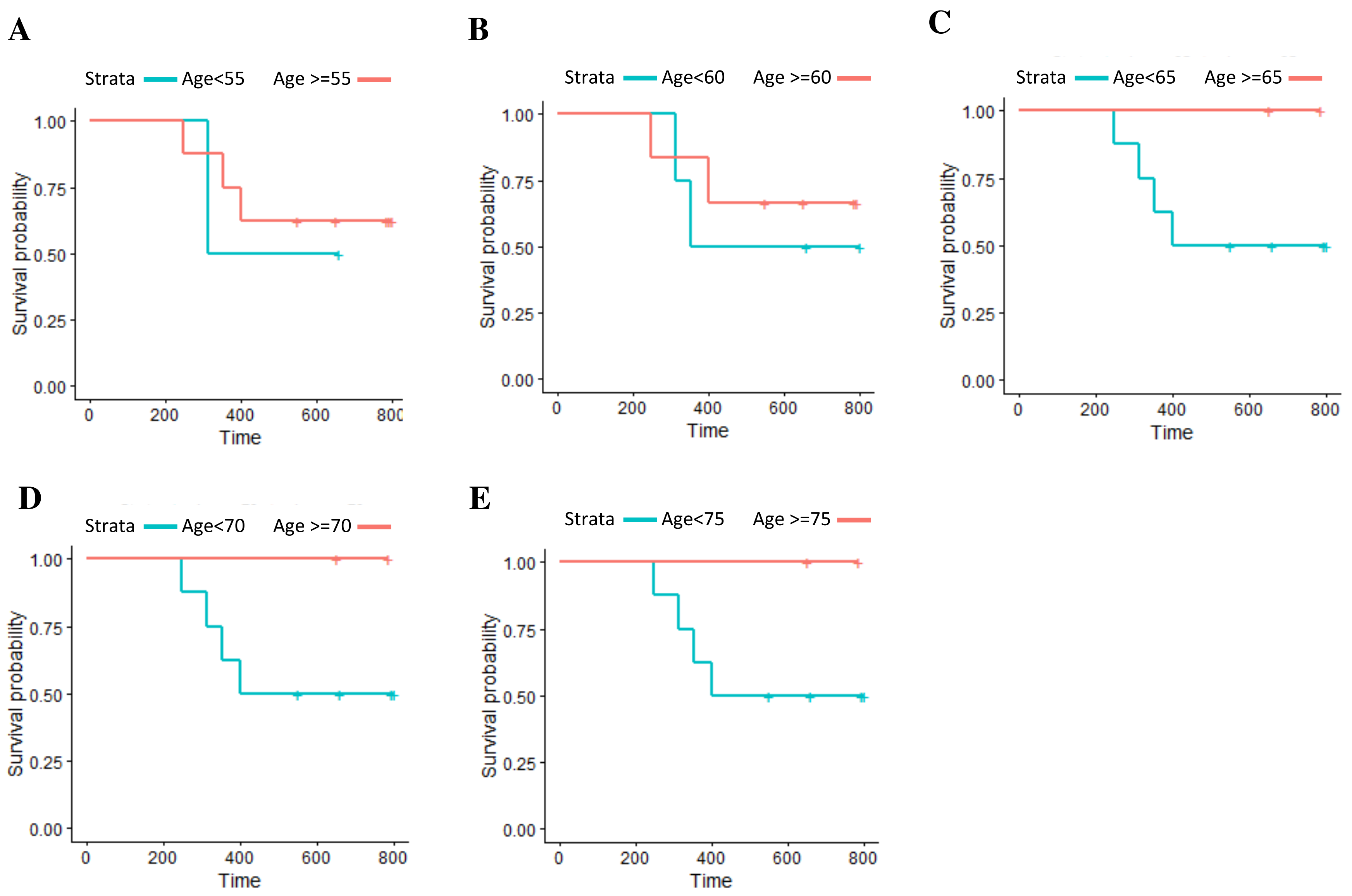

### Supplementary Figure 4

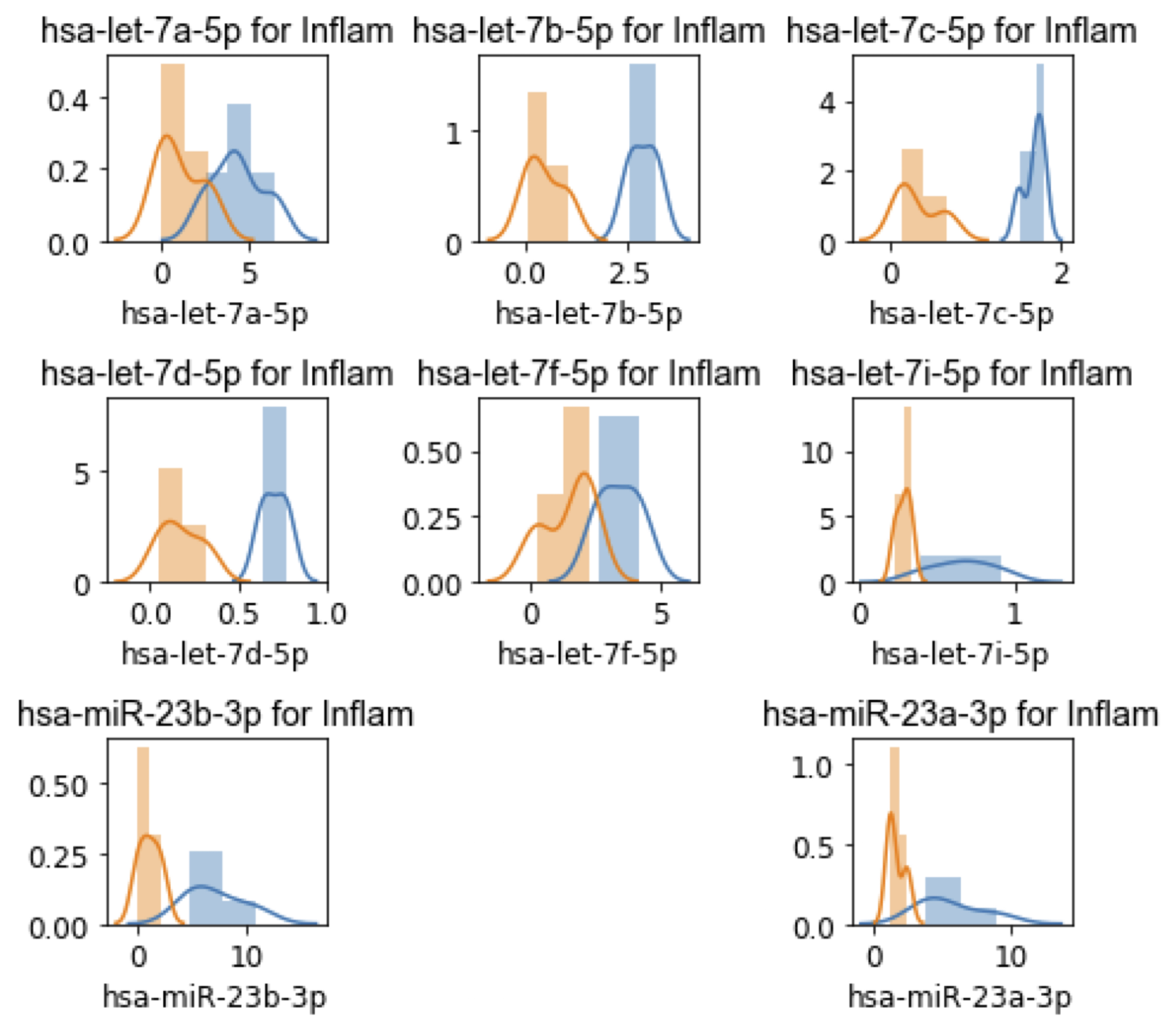
