## Supplementary Table 2 for "Inflammatory state of lymphatic vessels and miRNA profiles associated with relapse in ovarian cancer patients"

**LVs with high versus low inflammation**

| miRNA | Fold Regulation | p value |
|---|---|---|
| miR-301a-3p | 5.766 | 0.4688 |
| miR-144-3p | 4.173 | 0.1242 |
| miR-186-5p | 4.138 | 0.1288 |
| miR-19a-3p | 3.953 | 0.3073 |
| miR-19b-3p | 3.456 | 0.2669 |
| miR-101-3p | 3.075 | 0.2711 |
| miR-15a-5p | 3.011 | 0.2785 |
| miR-410-3p | 2.401 | 0.8776 |
| miR-497-5p | 2.319 | 0.2108 |
| miR-548e-3p | -2.064 | 0.4127 |
| miR-125a-5p | -2.200 | 0.1220 |
| **let-7i-5p** | **-2.233** | **0.0328** |
| let-7e-5p | -2.666 | 0.0802 |
| let-7g-5p | -2.986 | 0.0969 |
| **let-7f-5p** | **-3.198** | **0.0378** |
| **miR-23a-3p** | **-3.443** | **0.0401** |
| miR-545-3p | -3.473 | 0.2340 |
| miR-98-5p | -4.635 | 0.0823 |
| **let-7d-5p *** | **-5.057** | **0.0009** |
| miR-15b-5p | -5.238 | 0.0696 |
| miR-181d-5p | -5.972 | 0.1116 |
| miR-130a-3p | -6.144 | 0.7502 |
| **let-7c-5p *** | **-6.425** | **0.0003** |
| miR-607 | -7.332 | 0.1901 |
| miR-202-3p | -7.821 | 0.1220 |
| miR-656-3p | -8.224 | 0.2351 |
| **let-7b-5p *** | **-11.423** | **0.0006** |
| **miR-23b-3p** | **-15.166** | **0.0152** |
| **hsa-let-7a-5p** | **-188.204** | **0.0389** |

| up-regulated | down-regulated | **Bold= p < 0.05** |
|---|---|---|
