## Supplementary Table 3 for "Inflammatory state of lymphatic vessels and miRNA profiles associated with relapse in ovarian cancer patients"

| LVs with low versus [medium or high] inflammation | | |
|---|---|---|
| miRNA | Fold regulation | p-value |
| miR-410-3p | 10.146 | 0.442 |
| miR-144-3p | 4.022 | 0.104 |
| miR-301a-3p | 3.501 | 0.725 |
| miR-101-3p | 2.700 | 0.368 |
| miR-381-3p | 2.408 | 0.170 |
| miR-19a-3p | 2.306 | 0.472 |
| miR-497-5p | 2.168 | 0.168 |
| miR-19b-3p | 2.028 | 0.422 |
| miR-181c-5p | 2.019 | 0.186 |
| miR-186-5p | 2.000 | 0.408 |
| miR-15a-5p | 1.888 | 0.452 |
| miR-340-5p | -1.819 | 0.217 |
| miR-98-5p | -1.843 | 0.427 |
| miR-454-3p | -1.909 | 0.339 |
| miR-181d-5p | -1.938 | 0.987 |
| miR-130a-3p | -1.951 | 0.589 |
| **let-7i-5p** | **-2.021** | **0.013** |
| miR-374a-5p | -2.109 | 0.228 |
| miR-15b-5p | -2.154 | 0.268 |
| let-7d-5p | -2.291 | 0.566 |
| **let-7g-5p** | **-2.390** | **0.031** |
| miR-202-3p | -2.497 | 0.682 |
| miR-607 | -2.578 | 0.523 |
| miR-656-3p | -2.741 | 0.600 |
| miR-545-3p | -2.773 | 0.074 |
| let-7c-5p | -2.982 | 0.347 |
| miR-23b-3p | -3.654 | 0.406 |
| let-7b-5p | -4.443 | 0.336 |
| let-7a-5p | -14.817 | 0.793 |

up-regulated | down-regulated | **Bold= p < 0.05**
