## Supplementary Table 4 for "Inflammatory state of lymphatic vessels and miRNA profiles associated with relapse in ovarian cancer patients"

**Cancer-infiltrated vs non-cancer-infiltrated LVs**

| miRNA | Fold Regulation | p value |
|---|---|---|
| **miR-144-3p*** | **14.020** | **0.0010** |
| **miR-181c-5p** | **10.629** | **0.0170** |
| miR-301a-3p | 7.300 | 0.3090 |
| miR-19a-3p | 6.896 | 0.1970 |
| miR-19b-3p | 6.055 | 0.1470 |
| miR-101-3p | 5.083 | 0.0740 |
| **miR-381-3p** | **4.070** | **0.0140** |
| miR-181a-5p | 3.875 | 0.2740 |
| miR-130a-3p | 3.853 | 0.1150 |
| miR-186-5p | 3.214 | 0.1310 |
| let-7a-5p | 2.920 | 0.2870 |
| **miR-497-5p** | **2.902** | **0.0260** |
| miR-590-5p | 2.780 | 0.5210 |
| miR-15a-5p | 2.767 | 0.1820 |
| **miR-93-5p*** | **2.252** | **0.0010** |
| miR-29a-3p | 2.166 | 0.5250 |
| miR-29b-3p | 2.098 | 0.3200 |
| **miR-16-5p** | **2.077** | **0.0310** |
| **let-7i-5p** | **-2.080** | **0.0210** |
| let-7b-5p | -2.274 | 0.0700 |
| let-7e-5p | -2.505 | 0.4320 |
| let-7c-5p | -2.556 | 0.0780 |
| let-7d-5p | -2.672 | 0.1220 |
| let-7f-5p | -3.239 | 0.2700 |
| miR-202-3p | -4.800 | 0.2990 |

| up-regulated | down-regulated | **Bold= p < 0.05** |
|---|---|---|
