## Supplementary Table 5 for "Inflammatory state of lymphatic vessels and miRNA profiles associated with relapse in ovarian cancer patients"

**LVs from relapse versus non-relapse patients**

| miRNA | Fold regulation | p-value |
|---|---|---|
| miR-144-3p | 5.069 | 0.0906 |
| **miR-186-5p** | **4.930** | **0.0294** |
| miR-301a-3p | 3.750 | 0.4580 |
| miR-19b-3p | 3.267 | 0.2037 |
| miR-19a-3p | 3.183 | 0.2633 |
| miR-15a-5p | 2.975 | 0.1666 |
| miR-29b-3p | 2.609 | 0.1861 |
| miR-29c-3p | 2.475 | 0.4066 |
| miR-497-5p | 2.295 | 0.0672 |
| miR-101-3p | 2.117 | 0.2561 |
| miR-34a-5p | 2.089 | 0.0828 |
| miR-548d-3p | 2.009 | 0.6136 |
| miR-30e-5p | -1.827 | 0.3177 |
| miR-548e-3p | -1.917 | 0.2868 |
| let-7g-5p | -1.938 | 0.1780 |
| let-7f-5p | -1.954 | 0.4330 |
| miR-23a-3p | -2.769 | 0.1140 |
| miR-449a | -3.197 | 0.3953 |
| miR-15b-5p | -3.302 | 0.1104 |
| miR-98-5p | -3.430 | 0.0673 |
| let-7d-5p | -3.681 | 0.0645 |
| miR-181d-5p | -4.029 | 0.1767 |
| miR-130a-3p | -4.306 | 0.5101 |
| **let-7c-5p** | **-4.335** | **0.0385** |
| miR-301b-3p | -4.776 | 0.1515 |
| **let-7b-5p** | **-6.940** | **0.0383** |
| **miR-23b-3p** | **-8.529** | **0.0278** |
| miR-656-3p | -16.377 | 0.3143 |
| let-7a-5p | -42.416 | 0.2375 |

up-regulated | down-regulated | **Bold= p < 0.05**
