## Supplementary Figure 5 for "Inflammatory state of lymphatic vessels and miRNA profiles associated with relapse in ovarian cancer patients"

A

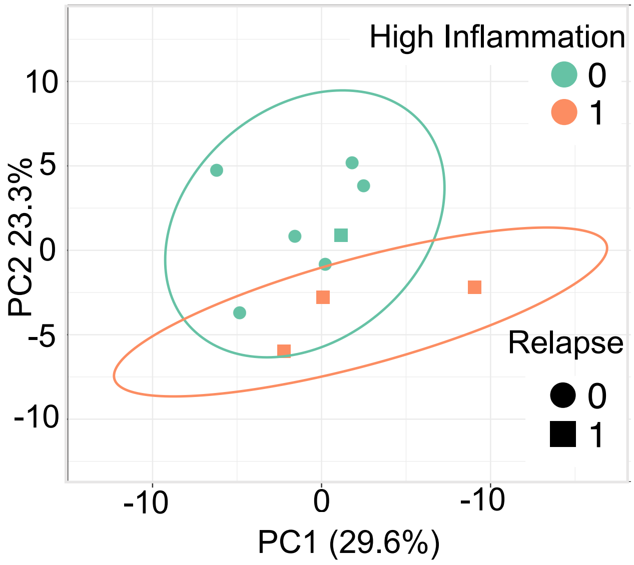


B

| PC1 | | PC2 | |
|---|---|---|---|
| **miR-29b-3p** | -0.222 | miR-125b-5p | 0.243 |
| miR-186-5p | -0.211 | miR-98-5p | 0.243 |
| **miR-301a-3p** | -0.210 | miR-125a-5p | 0.221 |
| **miR-101-3p** | -0.205 | miR-17-5p | 0.216 |
| **miR-19b-3p** | -0.195 | miR-30a-5p | 0.202 |
| **miR-181c-5p*** | -0.194 | **miR-23a-3p*** | 0.200 |
| **miR-19a-3p** | -0.192 | miR-15b-5p | 0.198 |
| miR-30b-5p | -0.192 | **let-7b-5p*** | 0.196 |
| **miR-497-5p*** | -0.191 | **let-7i-5p*** | 0.194 |
| miR-145-5p | -0.189 | miR-20b-5p | 0.188 |
| **miR-16-5p*** | -0.186 | **let-7d-5p*** | 0.187 |
| **miR-15a-5p** | -0.184 | miR-340-5p | 0.183 |
| miR-34a-5p | -0.171 | **let-7c-5p*** | 0.178 |
| miR-424-5p | -0.171 | **let-7a-5p*** | 0.171 |
| miR-20a-5p | -0.170 | miR-20a-5p | 0.168 |
| miR-128-3p | -0.165 | miR-545-3p | 0.165 |
| miR-30d-5p | -0.155 | **let-7f-5p*** | 0.151 |
| miR-29c-3p | -0.150 | miR-30e-5p | 0.149 |
| **let-7d-5p*** | 0.146 | miR-181b-5p | 0.147 |
| **let-7c-5p*** | 0.143 | **let-7e-5p*** | 0.145 |

C

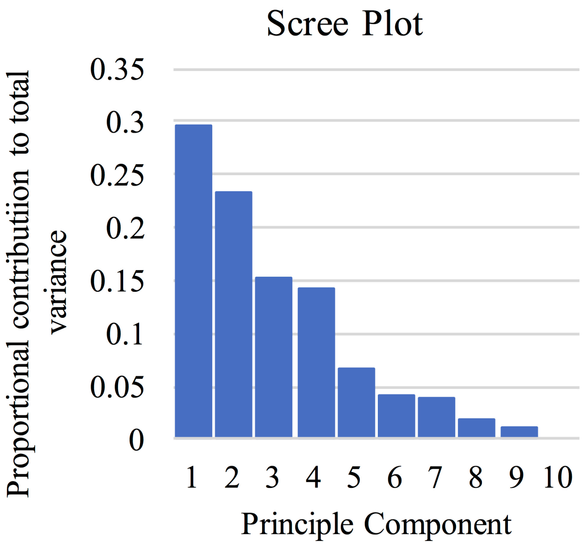
