## Supplementary Table 6 for "Inflammatory state of lymphatic vessels and miRNA profiles associated with relapse in ovarian cancer patients"

| Down-regulated miRNA | | | |
|---|---|---|---|
| KEGG pathway | p-value | Genes | miRNA |
| Tarbase | | | |
| Proteoglycans in cancer | 1.86E-11 | 88 | 3 |
| Viral carcinogenesis | 4.65E-08 | 77 | 3 |
| Fatty acid biosynthesis | 9.08E-08 | 4 | 3 |
| Cell cycle | 9.08E-08 | 58 | 3 |
| Lysine degradation | 1.42E-07 | 22 | 3 |
| Protein processing in endoplasmic reticulum | 1.56E-06 | 72 | 3 |
| Adherens junction | 1.64E-05 | 37 | 3 |
| p53 signaling pathway | 2.94E-05 | 36 | 3 |
| Epstein-Barr virus infection | 8.24E-05 | 84 | 3 |
| TGF-beta signaling pathway | 1.47E-04 | 33 | 3 |
| Hippo signaling pathway | 1.47E-04 | 55 | 3 |
| FoxO signaling pathway | 1.49E-04 | 58 | 3 |
| Chronic myeloid leukemia | 3.27E-04 | 33 | 3 |
| Focal adhesion | 1.12E-03 | 80 | 3 |
| TargetScan | | | |
| Mucin type O-Glycan biosynthesis | 1.78E-04 | 1 | 2 |
| Valine, leucine and isoleucine biosynthesis | 5.71E-04 | 1 | 2 |
| Signaling pathways regulating pluripotency of stem cells | 1.67E-02 | 6 | 2 |
| 2-Oxocarboxylic acid metabolism | 2.07E-02 | 1 | 2 |
| Valine, leucine and isoleucine degradation | 2.58E-02 | 2 | 3 |
| Biosynthesis of amino acids | 3.21E-02 | 2 | 2 |
| Micro-CT-DS | | | |
| ECM-receptor interaction | 3.00E-06 | 19 | 4 |
| Signaling pathways regulating pluripotency of stem cells | 3.00E-06 | 47 | 4 |
| TGF-beta signaling pathway | 3.20E-06 | 28 | 4 |
| ErbB signaling pathway | 1.32E-05 | 35 | 4 |
| Long-term potentiation | 1.85E-04 | 27 | 4 |
| Proteoglycans in cancer | 1.85E-04 | 58 | 4 |
| Mucin type O-Glycan biosynthesis | 3.89E-04 | 9 | 4 |
| Axon guidance | 4.95E-04 | 37 | 4 |
| mTOR signaling pathway | 9.62E-04 | 24 | 4 |
| Adrenergic signaling in cardiomyocytes | 1.03E-03 | 41 | 4 |
| FoxO signaling pathway | 1.05E-03 | 41 | 4 |
| Glutamatergic synapse | 1.31E-03 | 32 | 4 |
| Focal adhesion | 1.88E-03 | 57 | 4 |
