## Supplementary Table 7 for "Inflammatory state of lymphatic vessels and miRNA profiles associated with relapse in ovarian cancer patients"

| Method | Accuracy | Inflammation<br>McNemar's Test-pval | Cohen's Kappa | No Inflam | Yes Inflam |
|---|---|---|---|---|---|
| **Logistic Regression** | 40% | 0.61 | -0.25 | 25.0% | 50.0% |
| **K-Nearest Neighbours** | 50% | 1 | -0.04 | 75.0% | 33.3% |
| **Support Vector Machine** | 60% | 0.24 | 0.24 | 0.0% | 50.0% |
| **Random Forests Classifier** | 30% | 0.68 | 0.16 | 50.0% | 67.0% |
| **Gaussian Naive Bayes** | 50% | 0.07 | -0.04 | 0.0% | 83.3% |

| Method | Accuracy | Stage>=IV<br>McNemar's Test-pval | Cohen's Kappa | Stage <IV | Stage >=IV |
|---|---|---|---|---|---|
| **Logistic Regression** | 40% | 0.61 | -0.25 | 25.0% | 50.0% |
| **K-Nearest Neighbours** | 60% | 0.22 | 0.16 | 83.3% | 25.0% |
| **Support Vector Machine** | 60% | 0.04 | 0.16 | 100.0% | 0.0% |
| **Random Forests Classifier** | 30% | 1 | -0.45 | 33.3% | 25.0% |
| **Gaussian Naive Bayes** | 50% | 0.007 | -0.04 | 83.3% | 0.0% |
