## Supplementary Table 8 for "Inflammatory state of lymphatic vessels and miRNA profiles associated with relapse in ovarian cancer patients"

| Inflam | Method | Accuracy | Cohen's Kappa | No Inflam | Inflam | Mc Nemar p-value |
|---|---|---|---|---|---|---|
| **1-St.Sig miRNAs** | Logistic Regression | 60 | 0.17 | 0 | 100 | 0.04 |
| | K-Nearest Neighbours | 80 | 0.58 | 75 | 83.3 | 0.72 |
| | Support Vector Machines | 60 | 0.17 | 0 | 100 | 0.04 |
| | Random Forests | 70 | 0.37 | 50 | 83.3 | 0.45 |
| | Gaussian Naive Bayes | 80 | 0.58 | 75 | 83.3 | 0.72 |
| **2- St.Sig miRNAs** | Logistic Regression | 70 | 0.37 | 50 | 83.3 | 0.45 |
| | K-Nearest Neighbours | 80 | 0.58 | 50 | 100 | 0.29 |
| | SVM | 80 | 0.58 | 50 | 100 | 0.29 |
| | Random Forests | 70 | 0.37 | 50 | 83.3 | 0.45 |
| | Gaussian Naive Bayes | 80 | 0.58 | 75 | 83.3 | 0.72 |

| Relapse | Method | Accuracy | Cohen's Kappa | No Relapse | Relapse | Mc Nemar p-value |
|---|---|---|---|---|---|---|
| **2-St.Sig miRNAs** | Logistic Regression | 80 | 0.58 | 100 | 50 | 0.29 |
| | K-Nearest Neighbours | 90 | 0.79 | 100 | 75 | 0.5 |
| | Support Vector Machines | 80 | 0.58 | 83.3 | 75 | 0.72 |
| | Random Forests | 60 | 0.16 | 67 | 50 | 0.68 |
| | Gaussian Naive Bayes | 80 | 0.28 | 100 | 50 | 0.58 |
| **4-St.Sig miRNAs** | Logistic Regression | 70 | 0.37 | 83.3 | 50 | 0.45 |
| | K-Nearest Neighbours | 90 | 0.79 | 100 | 75 | 0.5 |
| | Support Vector Machines | 90 | 0.79 | 100 | 75 | 0.5 |
| | Random Forests | 60 | 0.16 | 67 | 50 | 0.68 |
| | Gaussian Naive Bayes | 80 | 0.28 | 100 | 50 | 0.58 |

| Cancer | Method | Accuracy | Cohen's Kappa | No Cancer | Cancer | Mc Nemar p-value |
|---|---|---|---|---|---|---|
| **2-St.Sig, miRNAs** | Logistic Regression | 60 | 0.17 | 100 | 0 | 0.04 |
| | K-Nearest Neighbours | 100 | 1 | 100 | 100 | 1 |
| | Support Vector Machine | 60 | 0.17 | 100 | 0 | 0.04 |
| | Random Forests | 100 | 1 | 100 | 100 | 1 |
| | Gaussian Naive Bayes | 100 | 1 | 100 | 100 | 1 |
| **3-St.Sig, miRNAs** | Logistic Regression | 60 | 0.17 | 100 | 0 | 0.04 |
| | K-Nearest Neighbours | 100 | 1 | 100 | 100 | 1 |
| | Support Vector Machines | 60 | 0.17 | 100 | 0 | 0.04 |
| | Random Forests | 100 | 1 | 100 | 100 | 1 |
| | Gaussian Naive Bayes | 100 | 1 | 100 | 100 | 1 |
| **5-St.Sig, miRNAs** | Logistic Regression | 60 | 0.17 | 100 | 0 | 0.04 |
| | K-Nearest Neighbours | 100 | 1 | 100 | 100 | 1 |
| | Support Vector Machines | 60 | 0.17 | 100 | 0 | 0.04 |
| | Random Forests | 100 | 1 | 100 | 100 | 1 |
| | Gaussian Naive Bayes | 100 | 1 | 100 | 100 | 1 |
| **6-St.Sig, miRNAs** | Logistic Regression | 70 | 0.37 | 83.3 | 50 | 0.45 |
| | K-Nearest Neighbours | 90 | 0.79 | 83 | 100 | 1 |
| | Support Vector Machines | 100 | 1 | 100 | 100 | 1 |
| | Random Forests | 100 | 1 | 100 | 100 | 1 |
| | Gaussian Naive Bayes | 100 | 1 | 100 | 100 | 1 |
| **7-St.Sig, miRNAs** | Logistic Regression | 80 | 0.58 | 100 | 50 | 0.45 |
| | K-Nearest Neighbours | 100 | 1 | 100 | 100 | 1 |
| | SVM | 100 | 1 | 100 | 100 | 1 |
| | Random Forests | 100 | 1 | 100 | 100 | 1 |
| | Gaussian Naive Bayes | 100 | 1 | 100 | 100 | 1 |
